# Conserved principles of spatial biology define tumor heterogeneity and response to immunotherapy

**DOI:** 10.1101/2024.10.18.619136

**Authors:** Vivek Behera, Alexander Guzzetta, Hannah Giba, Ue-Yu Pen, Anna Di Lello, Bipul Pandey, Benjamin A. Doran, Alessandra Esposito, Apameh Pezeshk, Aliya Husain, Christine M. Bestvina, Justin Kline, Marina C. Garassino, Arjun S. Raman

**Affiliations:** Department of Medicine, Section of Hematology/Oncology, University of Chicago, Chicago, IL, 60637; Duchossois Family Institute, University of Chicago, Chicago, IL, 60637; Department of Pathology, University of Chicago, Chicago, IL, 60637; Pritzker School of Molecular Engineering, University of Chicago, Chicago, IL, 60637; Center for the Physics of Evolving Systems, University of Chicago, Chicago, IL, 60637

## Abstract

The complexity of tumor microenvironments (TMEs) poses a substantial challenge to understanding tumor heterogeneity and clinical outcomes. By studying an ensemble of 262 diverse solid tumors, we uncovered a conserved, hierarchical architecture of transcriptionally covarying regions we term Spatial Groups (SGs). SGs corresponded to discrete biological units as benchmarked against multiple spatial technologies, and their nested organization revealed context-dependent constraints within tumors. Using SGs for comparing tumors, we derived a pantumor classification where immune spatial heterogeneity was the dominant axis of variation. This classification stratified response to immune checkpoint blockade in an out-of-sample cohort of non-small cell lung cancer patients. Statistical approximation techniques defined a sparse set of protein markers capturing system-level properties of TME spatial biology, demonstrating a framework for distilling genome-wide information into clinically deployable diagnostics. Our findings position the architecture of SGs as a general model unifying TME structure with biological function and clinical translation.

## Main

The tumor microenvironment (TME) is a complex milieu of interacting cells, proteins, and other biological components that influences critical properties of tumor biology such as growth, metastasis, and response to therapy^1,2^. Biological variation within the TME reflects clinically relevant differences across genetic, pathway, cellular, and tissue-level scales^3,4^. For instance, recent studies have demonstrated the prognostic and predictive power of TME-specific biomarkers, such as tumor infiltrating lymphocyte (TIL) score in melanoma and ‘Immunoscore’ – the spatial balance of CD3+ and CD8+ T cell density – in colorectal cancer^5–9^. These and other similar findings have motivated significant investment in studying the TME as an ecosystem of cells interacting within the spatial constraints of a tumor, most notably with technologies that couple cellular information about RNA or protein levels with cellular spatial locations^10^. Such spatial molecular profiling studies conducted in a variety of tumor types have revealed a common theme: the substantial heterogeneity within tumors (intratumoral) and across tumors (intertumoral) makes elucidating organizing principles of the TME very challenging^11^. By extension, the clinical utility of TME spatial profiling has been limited in scope.

Recent efforts have begun to outline a strategy for learning conserved aspects of TME spatial biology with the idea that these aspects reflect organizing principles that may be clinically relevant. These studies have collectively demonstrated the existence of recurrent multicellular spatial structures associated with tumor biology – somatic mutations, cell cycle synchrony, invasive fronts – and with cancer prognosis^12–18^. Obtaining these insights relied on imaging-based technologies that query tens of proteins to identify phenotypes such as cell type, cell cycle state, and a limited set of cell functional states. While these studies have been invaluable in demonstrating the relevance of spatial organization for TME biology, their reliance on limited sets of markers that were pre-selected based on prior knowledge precludes uncovering general principles of TME spatial organization that may emerge from a broader, unbiased approach. Spatial transcriptomics (‘ST-seq’) and related technologies, which provide genome-wide transcriptional information coupled with nearly single-cell resolution of spatial coordinates, enable a broad and unbiased assessment of TME spatial biology. However, the complexity of such data has hindered moving beyond mere description into an elucidation of spatial biology principles^19,20^. Advances in statistical inference developed in other fields of biology – protein science, genomics, and microbiome science – provide useful frameworks for addressing this challenge. For instance, at the scale of proteins, analysis of conserved amino acid covariation within ensembles of related proteins has yielded protein ‘sectors’ – groups of amino acids that are critical for engineering synthetically folded and functional proteins^21–24^. At the scale of genomes, covariation analysis of gene content across extant diversity within kingdoms of life has revealed units of collective protein-protein interactions that are critical for behavior and organismal fitness^25–28^. At the scale of microbiomes, covariation between bacterial taxa across individuals has yielded ‘ecogroups’ – groups of taxa that are of functional and clinical significance amongst humans^29–31^. Thus, these studies have established a general strategy for parsing organization amongst complex biological systems: first identify an ensemble of systems, then statistically deduce features that are conserved across the ensemble and critical for system function.

Using such studies as inspiration, we hypothesized that statistical analysis of ST-seq data across a diverse ensemble of solid tumors – a ‘pan-tumor’ database – may reveal conserved patterns of TME spatial biology in an unbiased manner. Here, we developed a new statistical formalism called TumorSPACE – a method that detects the statistical structure of covariation amongst cells or spots based on their profile of measured features (e.g., gene expression). Employing TumorSPACE across a database of 262 solid tumors that were profiled by ST-seq revealed the shared presence of hierarchically structured multicellular groups of transcriptionally covarying spots that we term Spatial Groups (SGs). We found SGs to be context-embedded spatial units, revealing nested domains of tumor microenvironments whereby a larger microenvironment influences the biology of smaller microenvironments within it. Using SGs as the basis for determining TME spatial domains, we defined a metric termed gene ‘spatial lability’ (SLAB) – a measure of the spatial heterogeneity of gene expression determined by expression differences across SGs. Aligning tumors by their genome-wide SLAB scores enabled comparative spatial transcriptomics and construction of a pan-tumor model of TME spatial biology. We found that the dominant axis of spatial variation across TMEs was related to immune biology. We therefore tested whether this classification could distinguish response to immune checkpoint blockade (ICB). Using an independent cohort of non-small cell lung cancer (NSCLC) patients, we found that classification by our pan-tumor spatial biology model accurately distinguished patients who responded to ICB from those who did not respond. Finally, we developed a simulation-based approach that leveraged tumor SGs to align genome-wide transcriptional data with mIF data from the same samples. We found that two separate panels of only seven IF markers were sufficient to recapitulate genome-wide spatial similarities observed between tumors.

Overall, our work describes a reparameterization of the TME into hierarchically structured units: Spatial Groups. This description unifies TME structural organization with biological context and clinical utility.

### Spatial Groups (SGs) define a conserved architecture of TME spatial biology

To identify general principles of TME organization, we assembled a multimodal database of 262 tumors spanning 18 tumor types (**Fig. 1A, Fig. S1, Supplementary Table 1**). All tumors were profiled by spatial transcriptomics (ST-seq, 10X Visium platform) and the paired hematoxylin and eosin (H&E)-stained images were annotated by a board-certified pathologist at the University of Chicago using a nested classification scheme that described properties of tumor tissue, tumor-adjacent normal tissue, and artifacts (**Fig. S2A**) (**Supplementary Table 2**) (Methods). We found that pathologist annotations were consistent with expectations of biological and technical variation amongst tumor types (**Fig. S2B-D**). We also performed 51-multiplexed immunofluorescence on eight tumors (mIF) using a panel designed to identify tumor cells, tumor immune cell subsets, and a limited set of functional cell states at single-cell resolution (Methods, **Supplementary Table 3**). These mIF data were annotated by the same pathologist using a nested classification scheme identifying cell types, ‘neighborhoods’ (local collections of particular cell types), and ‘sub-neighborhoods’ (smaller neighborhoods nested within neighborhoods) (**Fig. S3**) (**Supplementary Table 4**) (Methods).

**Figure 1.**
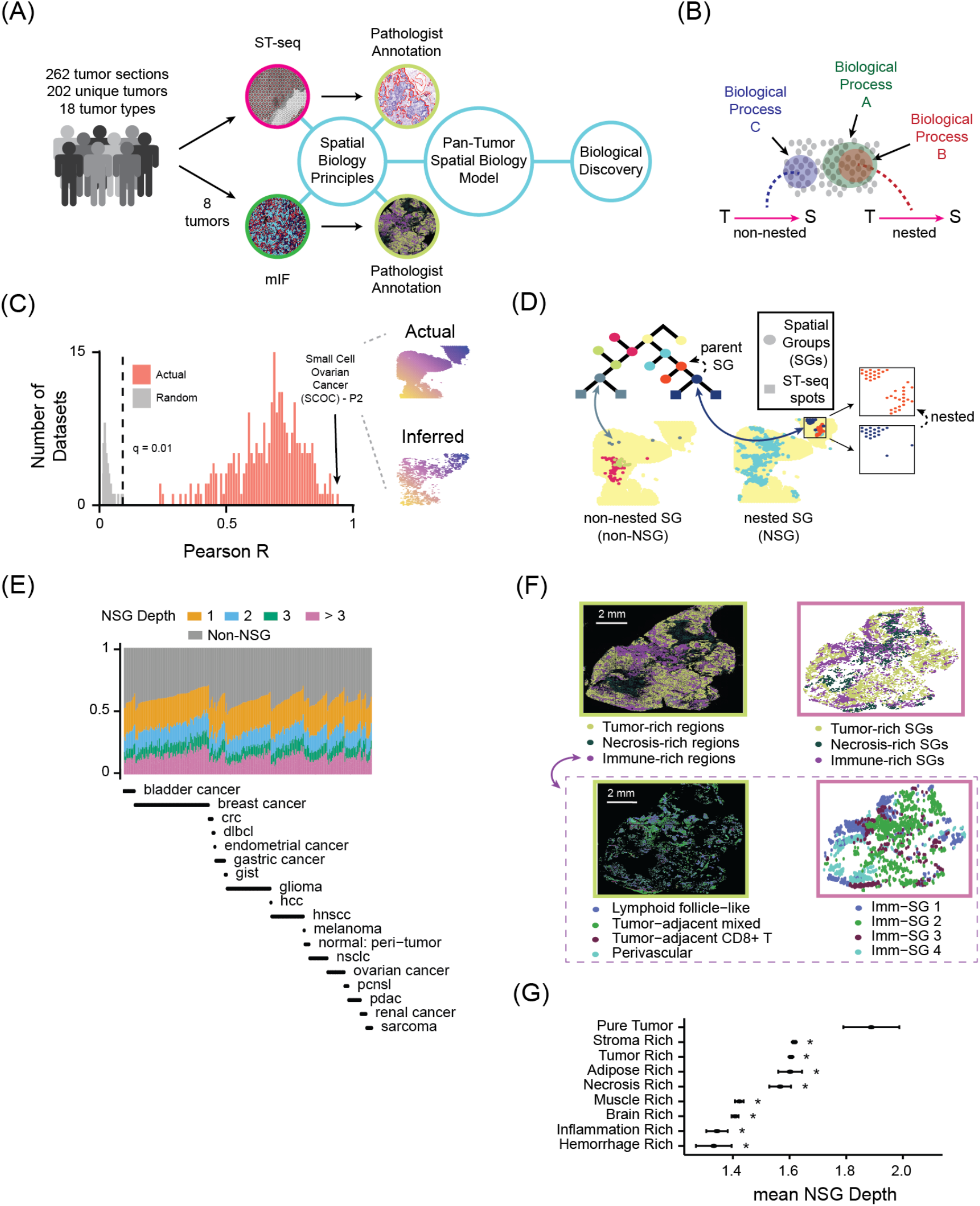
A conserved architecture of TME spatial biology. **(A)** As a strategy for identifying principles of TME spatial organization, we constructed a database comprised 262 tumors spanning 18 tumor types. Spatial transcriptional profiling and pathologist annotation was performed on all tumors. A subset of eight tumors were also profiled by 51-multiplexed immunofluorescence (mIF) and associated pathologist annotation. **(B)** A map (pink arrows) relating transcriptional (‘T’) and spatial (‘S’) information that captures non-nested and nested spatial contexts of biological processes. **(C)** Histogram of correlation values (Pearson R) between actual spatial distances for all pairs of spots within a tumor ST-seq dataset and pairwise distances inferred by TumorSPACE. Inset shows inferred spatial distribution of spot locations for the tumor exhibiting the highest Pearson R: small cell ovarian cancer sample ‘P2’ (SCOC-P2). **(D)** Relationship between TumorSPACE map and spatial distribution of spots in a sample. A part of the TumorSPACE map for a sample (SCOC-P2) is shown; squares are spots in the biopsy, each circle is a Spatial Group (SG). Highlighted are one non-nested and nested SG; inset illustrates nesting of spots in the blue SG within its ‘parent’ red SG. **(E)** Fraction of spots in an ST-seq dataset (y-axis) belonging to non-NSGs (gray bars) or NSGs of varying depth (colored bars) for all tumors in our database. X-axis is ordered and labeled according to tumor type. **(F)** Comparison of pathologist annotations (left) and SGs (right) that are aligned in spatial location at large (top) and intermediate (bottom) physical scales. **(G)** Mean NSG depth of spots (x-axis) grouped by pathologist-annotated tumor sub-region type (y-axis) across our tumor database. Non-NSGs are assigned a depth of 0. Error bars reflect the standard error of the mean. *Wilcoxon p-value < 0.05.

We first sought to construct a mapping that could infer TME spatial organization from ST-seq transcriptional data. Previous literature has demonstrated the presence and importance of spatially nested and non-nested biological processes in the TME^17,32^. As such, we wanted our mapping to distinguish nested and non-nested biological processes – a quality that currently developed frameworks for ST-seq data do not contain (**Fig. 1B**)^33–38^. We therefore developed a new framework called ‘TumorSPACE’ (<u>Tumor Sp</u>atial <u>A</u>rchitectures from the <u>C</u>omplete <u>E</u>igenspectrum) (Methods). TumorSPACE first uses patterns of transcriptional covariation to define hierarchical relationships between ST-seq spots^28,39^. This yields a tree-like model relating all spots in the TME where each leaf of the tree is an individual spot, branchpoints in the tree group spots together that are transcriptionally similar, and the root of the tree reflects the whole sample as a collection of spots. TumorSPACE then removes branches of that tree that do not relate to spatial organization (**Fig. S4**). The resulting model is a dataset-specific ‘TumorSPACE map’. We created TumorSPACE maps for all 262 datasets comprising our database. From these maps, we inferred spot locations either from TumorSPACE or by randomly matching them to locations in our dataset as a null model. For each dataset, we computed the correlation between spot-spot distances in the actual data versus inference. In each tested dataset, TumorSPACE maps inferred spot-spot distances significantly better than the null model (q < 0.01) (**Fig. 1C**) (Methods). Thus, TumorSPACE created accurate maps between transcriptional information and TME spatial organization.

Given this, we studied the collection of TumorSPACE maps to reveal any qualities of TME spatial organization that were conserved across our database. To understand the architecture of a TumorSPACE map, we first focused on the map exhibiting the highest Pearson correlation of inferred spot-spot distances with real spot-spot distances: a small-cell ovarian cancer dataset labeled ‘SCOC-P2’. We found that branchpoints in this map defined groups of spots that were spatially distributed in the biopsy sample. We therefore termed the branchpoints ‘Spatial Groups’ (SGs). In some cases, an SG was physically smaller – measured as the mean spot-to-spot distance within the group – than its ‘parent’ SG, the group of spots one layer closer to the root of the map (Methods). We referred to this phenomenon as ‘spatial nesting’, in line with terminology used by other groups that have found such behavior in specific tumor subsets^17,40^. We defined any ‘child’ SG that was spatially nested within its parent as a nested Spatial Group (NSG); any ‘child’ SG that was not spatially nested within its parent was a non-nested Spatial Group (non-NSG) (**Fig. 1D**) (Methods). Additionally, we found that in the SCOC-P2 dataset, parent NSGs could be spatially nested within grandparent NSGs, which in turn could be nested within great-grandparent NSGs and so on. To quantify this hierarchy, we defined ‘NSG depth’ as the number of NSGs encountered when moving from a given NSG toward the root of the TumorSPACE map before reaching a non-NSG (**Fig. S5**) (Methods). Systematic analysis of all tumors in our database revealed a spatial architecture of the TME that is broadly conserved: SGs are comprised of a consistent distribution of non-NSGs and NSGs that can be nested up to several degrees (**Fig. 1E**).

Manual annotation by a pathologist is the current standard approach for systematically identifying spatial domains that can be compared across tumors. We therefore sought to relate such expert-defined regions to our data-driven, statistically-defined units (SGs). To address this, we first examined tumors for which ST-seq and mIF were performed on serial sections. Six tumors had sufficient similarity between sections to align ST-seq and mIF spatial coordinates, enabling direct comparison of SGs with high-resolution pathologist annotations (**Fig. S6**) (Methods). As shown for a representative tumor, the largest SGs corresponded to tumor-rich, necrosis-rich, or immune-rich regions (**Fig. 1F**, top). Immune-related SGs further subdivided into four smaller SGs matching regions enriched for lymphoid-like follicles, tumor-adjacent CD8+ T cell clusters, perivascular immune niches, or mixed immune infiltrates (**Fig. 1F**, bottom). We next evaluated whether the nesting of SGs was related to pathologist-defined biological variation across tumors. Pathologist annotation revealed three dominant axes of variation: tissue category (e.g. tumor, normal, artifact), cellularity, and tissue subcategory (e.g. stroma-rich tumor, inflammation-rich tumor, normal pancreatic tissue, normal breast tissue). NSG depth was significantly associated with each axis and was 1) higher in tumor regions than in normal regions, 2) higher in areas of greater cellularity, and 3) higher in normal tissues composed of highly organized multicellular niches (**Fig. S7**). Within tumors, NSG depth was greatest in subregions expected to display broad heterogeneity in gene expression (e.g. pure tumor-rich) and lowest in regions with restricted expression programs (e.g. inflammation-rich or hemorrhage-rich) (**Fig. 1G**). Together, these data illustrated that SGs and their associated hierarchical architecture reflect TME spatial organization that is highly aligned with pathologist-defined domains, motivating a deeper investigation into the biology encoded by SGs.

### Spatial Groups reveal a nested hierarchy of biological domains across tumors

We developed a framework to relate SGs with TME biological processes in a high-resolution and unbiased manner. We computed three metrics – differential gene expression, differential cell type abundance, and differential pathway enrichment – across all pairs of ‘sibling’ SGs that arise from a common parent SG (**Fig. S8A**). For measuring differential cell type abundance, we inferred cell types within an SG using SpaCET, a deconvolution method for use on spot-level spatial transcriptional information. We found broad agreement between SpaCET-inferred cell types and expected cell types based on H&E-based pathologist annotations across our database of tumors (**Fig. S8B**). For measuring differential pathway enrichment, we applied overrepresentation analysis (ORA) to the differential genes found within each SG (Methods)^41^.

As a case study for relating SGs with TME spatial biology, we focused on one of the six NSCLC samples where we observed a clear spatial distinction from mIF data between two well-characterized biological states – tumor proliferation and tumor immune exhaustion (**Fig. 2A**, left)^9^. When we examined the TumorSPACE map for this tumor, we found that these two regions aligned with two Spatial Groups (SG13 and SG18) that emerged from a single branchpoint (SG12) (**Fig. 2A**, right). SG13 – aligned with the tumor proliferative region – was enriched for malignant cells, B cells, and multiple conventional dendritic cell subtypes (cDCs), while SG18 – aligned with the immune-exhausted region – was enriched for endothelial cells, macrophages, plasmacytoid dendritic cells (pDCs), and cancer-associated fibroblasts (CAFs) (**Fig. 2B**). As an independent validation of these SpaCET-inferred cell type measurements, we also found concordant shifts in cell populations using the mIF data directly (**Fig. S8C**). At the pathway level, SG13 was enriched for cell cycle, RNA processing, and membrane trafficking – pathways associated with tumor proliferation – while SG18 was enriched for signaling pathways such as RTK, GPCR, and cytokine signaling, as well as innate immune processes – pathways associated with immune exhaustion (**Fig. 2C**)^42^. Thus, this case study illustrated close alignment between SGs and well-studied, distinct biological states in the TME.

**Figure 2.**
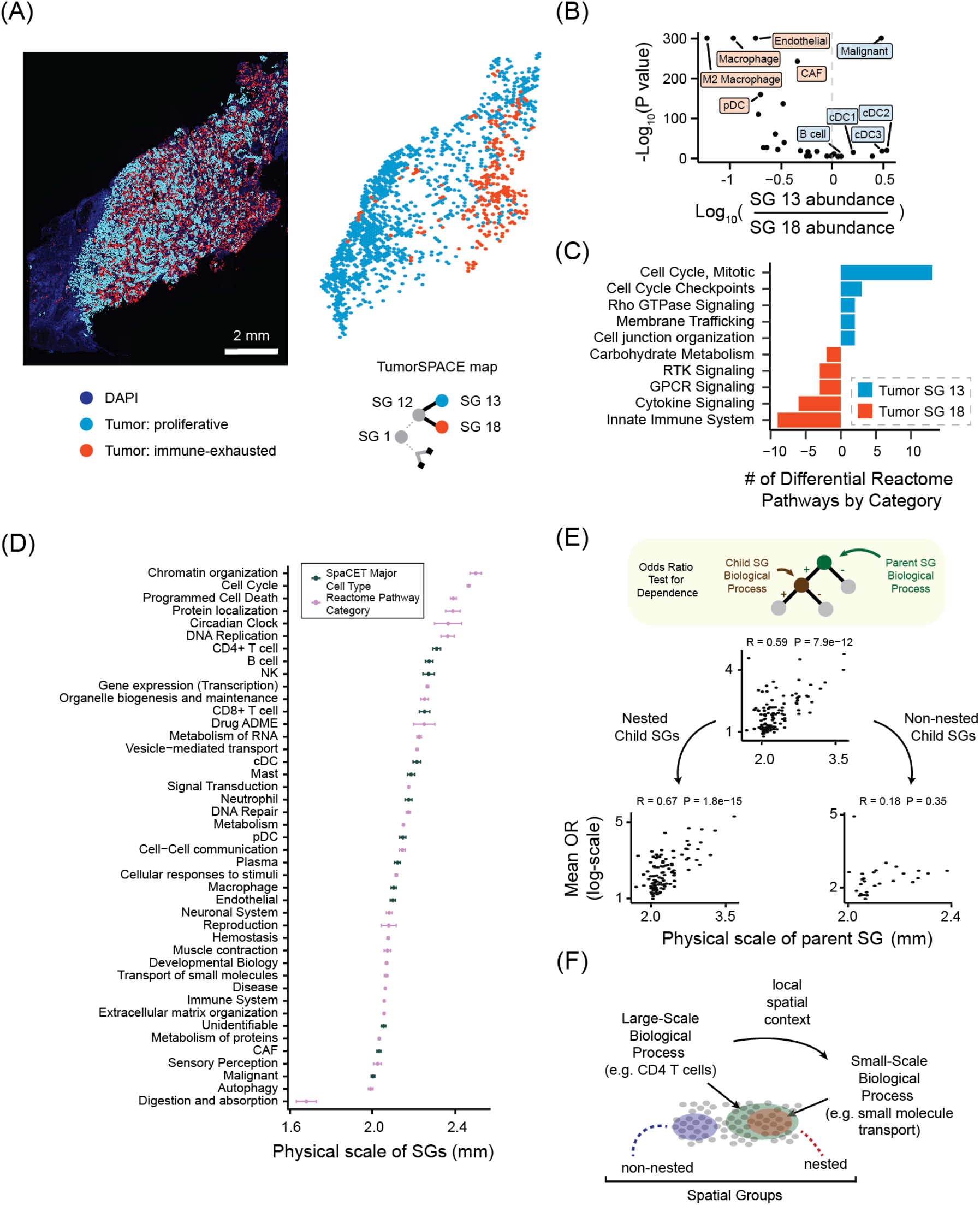
A model of TME spatial organization linking spatial scale with biological function. **(A)** Tumor sub-region identification either by multiplexed immunofluorescence (left) or the TumorSPACE map (right) for a representative NSCLC tumor. Fluorescent image shows DAPI (blue), proliferative tumor cells (cyan), and immune-exhausted tumor cells (red). TumorSPACE map shows spatial positions of spots colored by SG membership and hierarchical relationship between SG12 with SG13 and SG18. SGs are labeled such that the root of the TumorSPACE map is ‘SG1’. **(B)** Differential cell type abundance – shown as log10-transformed ratio of mean abundance (x-axis) and log10-transformed p-value (y-axis) – between SG13 and SG18 based on SpaCET deconvolution. Each point is a specific cell type. **(C)** Number of differentially enriched Reactome pathways between SG13 and SG18 (x-axis, see color key), grouped by pathway category (y-axis). **(D)** Physical scale of SGs as measured by mean of distances between pairs of spots (x-axis) for SGs differential in SpaCET-derived cell types (green) or pathway categories (pink). Analysis was performed for all SGs across all tumors in our database. Error bars reflect standard error of the mean. **(E)** (Top) Workflow for evaluating if a differentially abundant biological process within a parent SG influences a biological process within a child SG. (Middle, Bottom) Mean odds ratio (y-axis, absolute value, log2-transformed) of parent processes versus the physical scale of the parent SG (mm) where the child SG is nested or non-nested (middle), nested only (bottom left) or non-nested only (bottom right). Pearson correlation coefficients (R) and p-values are shown. **(F)** Model of TME spatial biology. TMEs are comprised of non-nested and nested Spatial Groups. Nested spatial groups encode large-scale processes that influence small-scale processes.

To move beyond individual case studies and assess whether SGs reflect generalizable patterns of biological organization across tumors, we performed a systematic analysis of all SGs across all TumorSPACE maps comprising our tumor database. For each SG, we computed the mean pairwise distance between all ST-seq spots within the SG – a proxy for its physical scale. We then examined how the physical scale of SGs varied across biological processes that were differentially enriched amongst all SGs. We found that specific processes were associated with SGs at characteristic physical scales in a consistent manner across tumors (**Fig. 2D**). Processes associated with large-scale SGs included, for example, CD4+ and CD8+ T cell abundance. These cell types have long been known to be excluded from large regions of tumors and were accurately identified to be spatially variable using early ST-seq analysis tools (e.g. SpatialDE) that considered the entire tumor to be a single domain^43,44^. Processes associated with intermediate-scale SGs included signal transduction, cell-cell communication, and chemokine responses.

These processes have previously been described as spatially variable when dividing tumors into ‘microregions’ on the scale of dozens of cells^45^. Finally, the processes associated with the smallest-scale SGs were often related to local tumor-immune interactions, a subset of TME biology that has been identified as spatially variable using tools such as SpaceMarkers that focus on local cell-cell interaction fronts^46^. Thus, our results illustrated that SGs collectively reflect differential biological processes spanning from tumor-wide physical scales to local spatial niches.

We next sought to interrogate how differential biological processes spanning many physical scales were related to each other. One possibility was that these processes were largely independent from each other and therefore biological context would not be set by a larger parent SG for a smaller child SG. Alternatively, prior work has shown examples of biological processes that exist within a nested hierarchy where processes acting within large spatial domains (i.e., hypoxia) set a strong context for processes within smaller domains (i.e., tumor-immune cell interactions)^17,32,47^. In this model, large parent SGs might set the biological context for smaller child SGs that they contain. To distinguish between these possibilities, we developed a permutation-based odds ratio (OR) test for biological dependence amongst differential pathways and cell types between all SGs in our tumor database and their associated child SGs (**Fig. 2E**, upper) (Methods). We found that larger physical scale spatial domains set a strong spatial context while smaller domains did not, but this result was only observed for NSGs and not non-NSGs (**Fig. 2E**, middle and lower).

Together, our findings spanning **Figs. 1** and **2** supported a model of the TME as a nested hierarchy of spatially organized biological processes (**Fig. 2F**). NSGs encode biological programs that are spatially embedded within larger SGs and thereby influenced by the broader spatial and functional context in which they reside. In contrast, non-NSGs encode biological programs that may be less dependent on their surrounding spatial environment. The architecture of SGs thus provides a data-driven, unbiased, and broadly conserved model linking spatial structure with biological function in the TME. This motivated evaluating whether SGs would be a useful unit of comparison across diverse tumors.

To compare the spatial biology of tumors, it is necessary to standardize descriptions of tumors in a way that retains important spatial information. Thus, we defined a compression of differential gene expression across SGs that we term gene ‘spatial lability’ (SLAB). This metric captures the extent to which gene expression is spatially heterogeneous across the whole TME. The process for defining gene SLAB was as follows. We first identified all SGs for a given TME. Then, for a given gene, we identified the SG siblings between which the gene was differentially expressed (**Fig. 3A**, left). To maximize the dynamic range of SLAB as a metric, we chose to define the smaller of the two sibling SGs, by number of constituent spots, as the SG reflecting a change in expression (**Fig. 3A**, middle). Finally, we computed the fraction of ST-seq spots represented by the union of these SGs relative to all ST-seq spots in the tumor. We termed this fraction the SLAB score for the gene of interest (**Fig. 3A**, right) (Methods).

**Figure 3.**
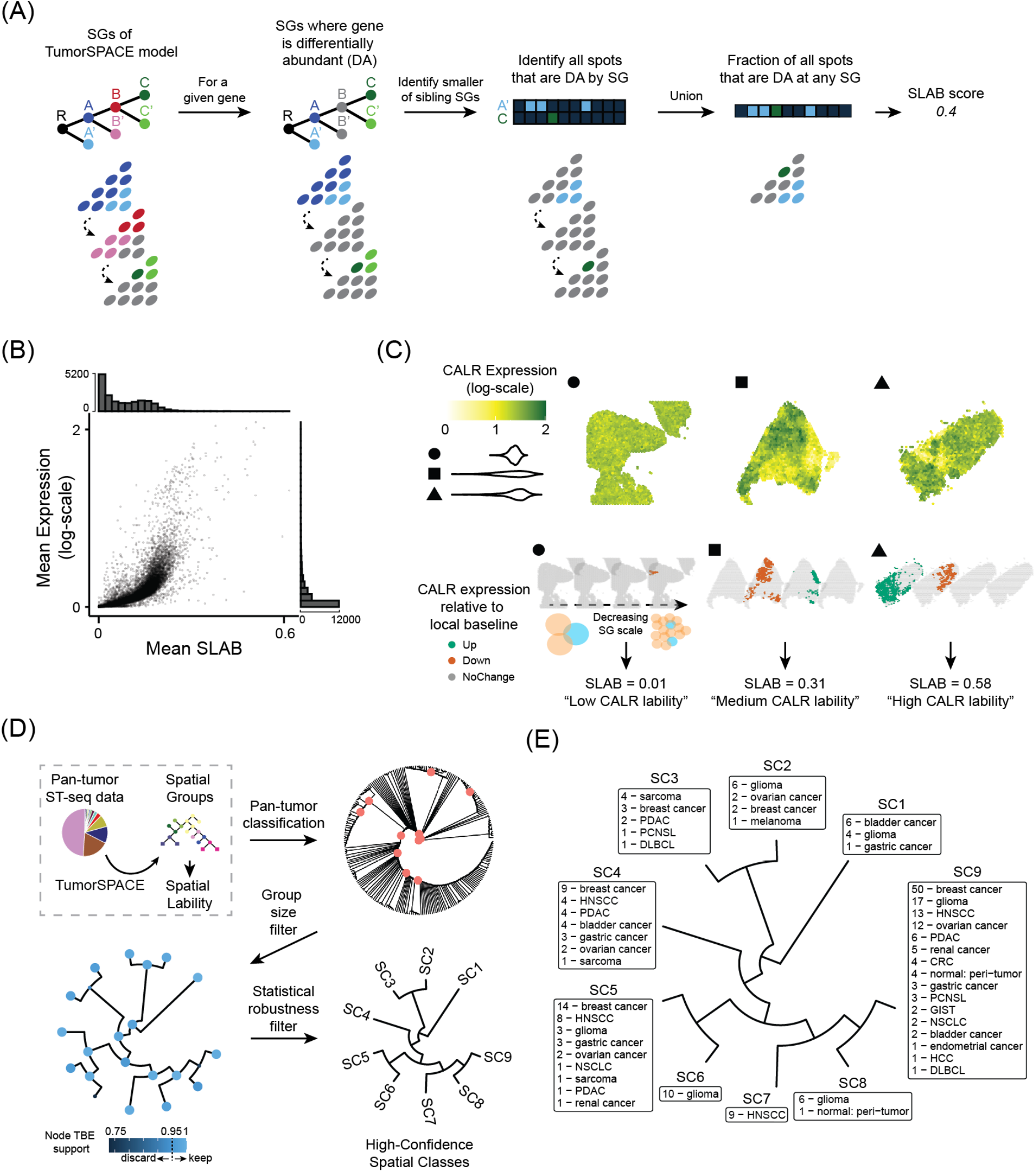
A spatially-aware pan-tumor classification of TMEs. **(A)** Schematic for computing Spatial Lability (SLAB) score for a given gene. First, all pairs of sibling SGs are identified where the gene is differentially expressed between sibling SGs. Second, for each pair of sibling SGs, the SG with fewer spots is identified. Third, the union of spots within these SGs is computed across the entire sample. The SLAB score is the fraction of all spots in the sample represented by step three. **(B)** Mean expression (log-scale, y-axis) versus mean SLAB score (x-axis) for all genes (dots) in our pan-tumor database. Histograms on each axis reflect the number of genes. **(C)** Upper panel shows spatial distribution of calreticulin (CALR) expression across three different samples (circle, square, and triangle) where average CALR expression is the same (see **Fig. S9B**). Lower panel shows distribution of spots within SGs that are differential in CALR expression for each sample as the physical scale of SGs decreases from left to right. Spots are colored by direction of expression change (colorkey) relative to local baseline. Below each depiction of decreasing SG scale, CALR SLAB score and associated interpretation (‘Low’, ‘Medium’, or ‘High’ CALR spatial lability) is shown. **(D)** Workflow for defining tumor Spatial Classes. Following steps in **Fig. S9C**, a pan-tumor classification tree is created, then filtered by group size. The tree is then filtered for statistical robustness to yield the final tree defining Spatial Classes. **(E).** Distribution of tumor types for all Spatial Classes. Numbers next to each tumor represent the number of samples within our pan-tumor cohort.

A comparison of SLAB scores with gene expression for all genes across all tumors in our database illustrated that SLAB scores were positively correlated with average gene expression but also captured other modes of gene-level spatial variation. For example, genes with low average expression exhibited variation in SLAB score (**Fig. 3B**). Furthermore, 47% of genes had no correlation between bulk gene expression and SLAB score when comparing across tumors (**Fig. S9A**). As a specific example, the calreticulin gene (CALR) had similar bulk expression levels in 3 selected tumors, yet its SLAB score varied from high to low across these tumors (**Fig. S9B**). Visualization of CALR expression across the TME clearly illustrated that the degree of spatial heterogeneity in gene expression followed the degree of spatial lability across tumors, as measured by SLAB score (**Fig. 3C**, top). Examination of the SGs associated with differential expression of CALR showed that the tumor with a high CALR SLAB score contained changes in CALR expression at both large- and medium-sized SGs, while the tumors with lower CALR SLAB scores had SG changes restricted to medium and small SGs (**Fig. 3C**, bottom). Together, these data demonstrate that SLAB scores capture information about the spatial heterogeneity of gene expression across the TME that is distinct from bulk gene expression.

We next sought to leverage SLAB, a standardized description of tumor spatial biology, to compare tumors to one another. To this end, we divided our tumor database into two cohorts: (i) a ‘pan-tumor cohort’ comprising 246 tumor sections across 18 tumor types and (ii) a single-institution cohort of 16 non-small cell lung cancer (NSCLC) tumors. The NSCLC cohort, obtained from the University of Chicago thoracic biobank, was unique in having matched retrospective clinical metadata about disease course and treatment outcomes. We reasoned that partitioning our database in this way could test whether a pan-tumor model of SG biology offered forward insights about clinical outcomes in an independent, out-of-sample collection of tumors. Because SLAB scores can be computed for every gene in the genome, the profile of genome-wide SLAB scores is a quantitative representation of genome-based spatial organization for each tumor. We therefore constructed a matrix where each row corresponded to a tumor within the pan-tumor cohort, each column to a gene, and each entry to the SLAB score of a gene in a tumor (**Fig. 3D**, top left; **Fig. S9C**, left). Pairwise Euclidean distances between tumors were computed and hierarchical clustering of the resulting distance matrix yielded a classification tree in which tumors clustered together based on similarity in their genome-wide SLAB profiles (**Fig. 3D**, top right; **Fig. 9C**, right). To ensure robustness of class assignments, we filtered for groups containing at least five tumors and performed bootstrap resampling that retained only branches with high statistical support (**Fig. 3D**, bottom left) (Methods).

The resulting classification yielded nine ‘Spatial Classes’ that we termed SC1 through SC9 (**Fig. 3D**, bottom right). Across our pan-tumor cohort, we found that 55% of tumor replicate pairs were classified within the identical SC, despite the majority of pairs coming from non-adjacent replicate sections within a tumor (**Fig. S10A**). As Spatial Classes were identified considering our whole pan-tumor cohort, it remained unknown whether smaller ST-seq cohorts could define these classes due to limitations in finite sampling and potentially rare abundance of certain Spatial Classes. We therefore performed a simulation with the following steps (Methods). First, tumors were added to the simulation one at a time in a random fashion; this defined an ‘epoch’ of the simulation where epoch 1 contained a single tumor, epoch 10 contained ten tumors, and so on. For each epoch, we estimated simulated Spatial Classes and compared these to the true Spatial Class designations as defined by the entire model. We did this by using an F1 statistic: higher mean F1 scores indicated that the true Spatial Class designation could be recovered with greater precision and recall across simulations. In running 100 such simulations, we found that certain Spatial Classes (SC1, SC4, and SC9) could be consistently defined with a small amount of data; one (SC8) required nearly the entire pan-tumor cohort for identification; several (SCs 2, 3, 5, 6, and 7) steadily increased performance as tumors were added to the training corpus (**Fig. S10B**). Finally, we noted an additional potential source of technical variability in computing the Spatial Class for any tumor. TumorSPACE computes for any tumor a collection of top-scoring models ranked by their ability to predict ST-seq spot locations. Thus, it is not guaranteed that the best and second-best TumorSPACE models for a given tumor have the same genome-wide SLAB scores or estimated Spatial Class. To test this, we first selected three pan-tumor datasets ranging in TumorSPACE model performance and identified the top five ranking models for each dataset (**Fig. S10C**). We then computed genome-wide SLAB scores for all of these ‘near-optimal’ models and estimated their Spatial Classes. Doing so demonstrated that all but one of these near-optimal models had the same estimated Spatial Class as their top-ranked ‘optimal’ model across the three datasets (**Fig. S10D**). Together, these data demonstrate Spatial Classes to be robust descriptions of tumor spatial biology that benefit from collecting ST-seq data at the scale of our pan-tumor cohort for their identification.

Analysis of the resulting Spatial Classes revealed that three classes were enriched for individual tumor types: SC6 and SC7 were composed entirely of gliomas and head and neck squamous cell carcinomas (HNSCC) respectively; SC8 consisted of gliomas and a biopsy sample originally labeled as ‘normal; peri-tumor’ that was re-classified as a glioma in pathology re-review. However, all other Spatial Classes were comprised of tumors from diverse tumor types **(Fig. 3E**). These findings therefore encapsulated two contrasting ideas: some studies have shown evidence for tumor type-specific spatial biology while others have demonstrated conserved properties of spatial biology across different tumor types^48^. We reasoned that by investigating the biological properties of our classification, we could learn how tumor type-specific and tumor type-agnostic factors fit together into a more unified understanding of TME spatial biology.

### Biological programs defining Spatial Classes

To approach understanding the biology distinguishing Spatial Classes in an unbiased manner, we evaluated the differences in SLAB profiles at the branchpoints in our pan-tumor classification (**Fig. 4A**, left) (**Supplementary Table 5**). We then applied ORA pathway analysis to identify biological programs enriched in each branch (**Fig. 4A**, right) (**Supplementary Table 6**). Hierarchical clustering of enriched pathways revealed that distinct branchpoints were dominated by a small number of biological axes: adaptive immunity, innate immunity, neurodevelopment, and metabolism (**Fig. 4B**). To confirm assignment of branchpoints to these biological axes, we investigated the differential genes and pathways that were driving classification. The three branchpoints highlighting differences in adaptive immunity exhibited the highest differential SLAB in genes related to 1) T cell chromatin remodeling through the NuRD (GATA2DA, MBD2) or SWI/SNF (ARID2) complexes, 2) lymphocyte development (TCF20, LIF4), or 3) intratumoral T cell function (SNX19, ZNF671) (**Fig. 4C**, upper left)^49–55^. The branchpoint corresponding to innate immunity SLAB variation was notable for largely unstudied genes (MAGEB17, ZNF773, DERPC) and pathways related to either general immune function (GPCR and cytokine signaling) or innate immunity specifically (**Fig. 4C**, upper right).

**Figure 4.**
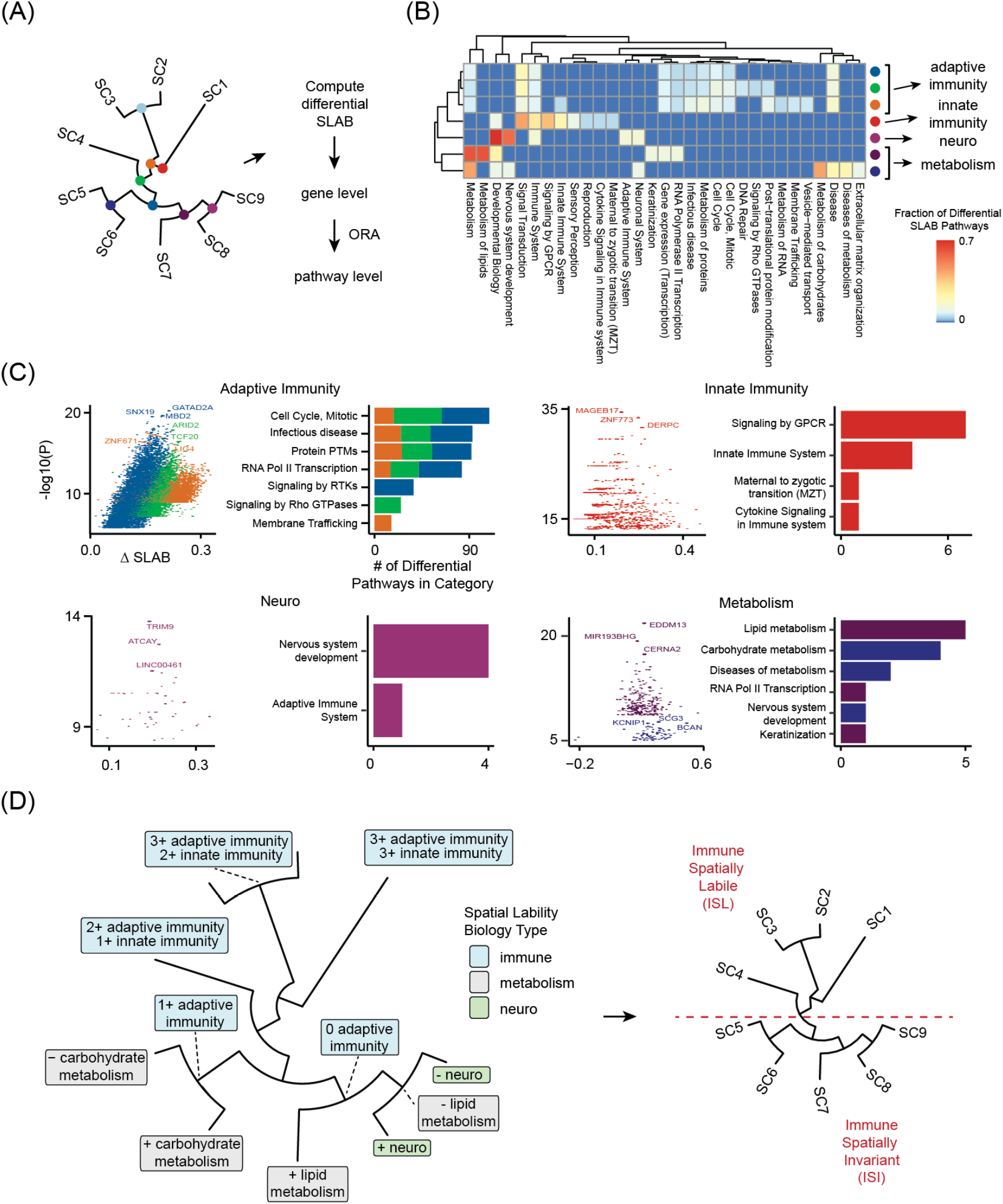
Spatial Classes reveal biological axes of tumor spatial biology. (**A)** Workflow for evaluating gene and pathway SLAB differences at all branchpoints distinguishing Spatial Classes. **(B)** Heatmap showing the fraction of differential SLAB pathways (color key) within a Reactome pathway category (columns) for each Spatial Class branchpoint (rows). Heatmap is hierarchically clustered, revealing four biological categories. **(C)** Interrogation of genes and pathways that exhibit statistically significant variation in SLAB for each of the biological categories shown in panel B **(C). (D)** Pan-tumor classification where branchpoints are labeled by biology that is spatially labile per categories shown in panel B (left). Red line distinguishes Spatial Classes by spatial lability in immune biology: ‘Immune Spatially Labile’ (ISL) versus ‘Immune Spatially Invariant’ (ISI).

Nested within immune function as the primary axis of SLAB variation was a secondary axis: metabolism. The branchpoint that distinguished SC6 – a group comprised of only gliomas – from SC5 was notable for variation in carbohydrate metabolism, with the most significant genes (KCNIP1, SCG3, BCAN) being closely related to glioma biology (**Fig. 4C**, bottom right)^56–58^. These data are consistent with reports that carbohydrate metabolic reprogramming – glycolysis versus oxidative phosphorylation – is directly linked to spatial organization in these tumors^17,59^. The other metabolism-oriented branchpoint distinguished SC7 – a group of purely HNSCC cancers – from SC8/9 and highlighted significant variation in lipid metabolism, including the top miRNA gene MIR193BHG, which has been shown to regulate cholesterol metabolism in a hypoxia-inducible manner^60^. Furthermore, lipid metabolism reprogramming has been demonstrated to be a key factor in HNSCC tumor growth and therapeutic resistance^61,62^.

The deepest axis of SLAB variation that we identified was in neurodevelopmental biology. This axis was identifiable only in the branchpoint distinguishing SC8 – another group comprised of only gliomas – from SC9 and was defined by neuron-restricted genes such as TRIM9 and ATCAY (**Fig. 4C**, bottom left)^63,64^.

Together, our results demonstrated a hierarchy of biological axes that describe variation in tumor spatial organization (**Fig. 4D**, left). Immune function was the dominant axis in this hierarchy, consistent with other findings of tumor immune biology having consistent effects on spatial organization across tumor types^20^. Based on this result, we defined Spatial Classes 1 through 4 as ‘Immune Spatially Labile’ (ISL) and 5 through 9 as ‘Immune Spatially Invariant’ (ISI) (**Fig. 4D**, right). We hypothesized that stratifying tumors by these categories might correlate with therapeutic response to ICB. We therefore turned to our NSCLC cohort for whom we had retrospective clinical data on response to ICB-containing treatment regimens.

### Using our classification of TMEs to predict therapeutic response

Despite substantial improvements in overall survival with the use of ICB in the metastatic NSCLC frontline setting, 5-year overall survival remains quite poor at 19%^65^. Moreover, the only clinically approved biomarker of response to ICB therapy – PD-L1 immunohistochemistry (IHC) – is weakly predictive of outcomes in the frontline metastatic setting for NSCLC, prompting ongoing studies on whether gene expression or cell type abundance biomarkers might be more predictive of such outcomes^4,65–67^.

Patients were selected to be in our NSCLC cohort based on having received frontline immune checkpoint blockade (ICB) immunotherapy with or without chemotherapy (Methods). For each sample, we computed genome-wide SLAB profiles and categorized them by their immune spatial lability as per the pan-tumor classification scheme defined in **Fig. 3E** (Methods). We then compared progression-free survival (PFS) of the 16 patients as defined by classification of immune spatial lability (ISL versus ISI) versus the clinical standard of PD-L1 IHC (**Fig. 5A**). In doing so, our strategy evaluated whether the pan-tumor classification could predict ICB response in the NSCLC cohort. Here, we emphasize that we use the term ‘prediction’ in the sense of a forward statistical test: no information from our NSCLC cohort was used to develop the pan-tumor definitions of ISL and ISI. Although larger studies would be required for establishing ISL/ISI as a predictive clinical biomarker, our study was designed to evaluate whether such a biomarker might be effective in stratifying ICB response^68^. Two possible variables were identified that could confound an association with ICB response: ICB regimen choice and presence of the somatic mutation KRAS G12C, which is targetable in the second line (**Fig. S11 A,B**). Univariate analysis found that neither variable was associated with PFS, excluding the possibility that these factors influenced our study (**Fig. S11C**).

**Figure 5.**
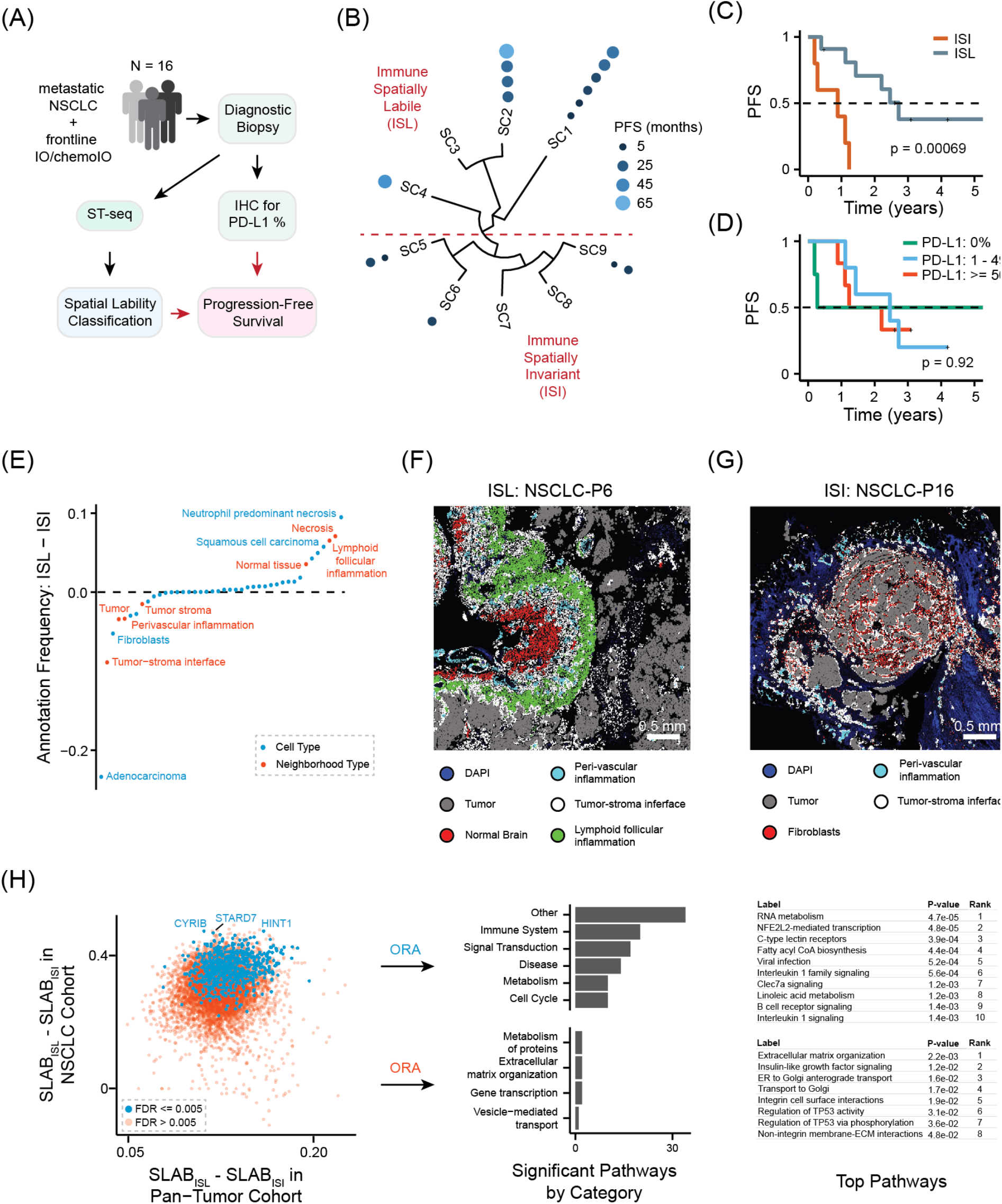
Pan-tumor classification distinguishes responders to immune checkpoint blockade in metastatic non-small cell lung cancer (NSCLC). **(A)** Workflow for evaluating progression-free survival (PFS) using either PD-L1 immunohistochemistry or spatial lability classification in our retrospective cohort of patients with metastatic NSCLC receiving frontline immunotherapy (IO) or IO plus chemotherapy. **(B)** Projection of the NSCLC cohort onto pan-tumor Spatial Classes. Each dot is a NSCLC sample, and the size and color describe progression-free survival (PFS, see color key). **(C,D)** Kaplan-Meier survival curves for categorizing the NSCLC cohort by ISL versus ISI (**C**) or PD-L1 status (**D**). **(E)** Comparison of cell type frequencies and coarse cellular neighborhood frequencies as determined by pathologist annotation between the ISL and ISI groupings. Features are ordered on the x-axis by rank on the y-axis. **(F)** Representative images of an ISL (**F**) and ISI (**G**) NSCLC tumor where specific biological features are identified by multiplexed immunofluorescence (see color key). **(H)** Differential SLAB analysis between ISL and ISI in the NSCLC cohort (y-axis) versus the pan-tumor cohort (x-axis). Overrepresentation analysis (ORA) defines pathways that are selectively differential in the ISL versus ISI categorization within the NSCLC cohort (right, top panel) relative to the pan-tumor cohort (right, bottom panel).

Of the 16 NSCLC samples, 11 were classified into Spatial Classes belonging to the ISL category (SC1, SC2, and SC4) while five samples were classified into Spatial Classes belonging to ISI (SC5, SC6, and SC9) (**Fig. 5B**). Strikingly, patients whose tumors fell into the ISL category exhibited significantly longer PFS than those in the ISI category (p = 0.00069) (**Fig. 5C**). In contrast, when patients were stratified using standard PD-L1 IHC thresholds (0%, 1-49%, and greater than or equal to 50%), no significant differences in PFS were observed (p = 0.92), nor were differences detected when applying a binary PD-L1 cutoff of 50% (p = 0.99) (**Fig. 5D, Fig. S12A**). Multivariate analysis of ISL/ISI in combination with a second covariate – 1) ICB regimen choice, 2) PDL1 status, 3) the presence of a KRAS G12 mutation, 4) the presence of an STK11 mutation, or 5) the presence of either an EGFR or BRAF mutation – demonstrated in each case that immune spatial lability was significantly predictive of PFS while the second covariate was not (**Fig. S12B,C**). Moreover, eight out of the twelve patients with measurable disease at treatment onset demonstrated shrinkage in tumor volumes shortly after treatment began, suggesting that classification by PFS was detecting differences in treatment response and durability rather than in treatment-agnostic factors such as disease prognosis (**Fig. S12D**) (Methods).

We next investigated whether the predictive capacity of the ISL/ISI classification might be attributable to sparse biological processes (e.g. CD8+ T cell abundance or a T cell exhaustion gene set) identifiable from bulk gene expression data. Using both our pan-tumor and NSCLC cohorts, we created sample-level bulk estimates of 1) SpaCET-deconvoluted cell type abundances and 2) expression of genes found within eight published gene set signatures of ICB response in NSCLC, including T cell effector function, T cell exhaustion, and TGF-*β* signaling (**Supplementary Table 7**) (Methods)^69–75^. As a comparison, we also generated SLAB scores corresponding to each cell type and gene signature. We found that none of these sparse biological processes could predict ICB response as well as our ISL/ISI classifier. Furthermore, eight of the top 10 results came from SLAB representation of these processes rather than bulk estimates (**Fig. S13A**). For example, the most predictive process – endothelial cell abundance – was only significant when represented as tumor-wide SLAB scores (p = 0.011) but not when represented as bulk abundance (p = 0.61) (**Fig. S13B,C**). On the other hand, the best bulk-estimated processes for stratifying ICB response in our NSCLC cohort were SpaCET-inferred plasma cell abundance and a B-cell gene signature, consistent with data that plasma cell bulk estimates can stratify ICB response in large NSCLC cohorts^76^.

To assess whether ISL and ISI distinctions reflect known features of ICB response, we compared these classifications to high-resolution pathologist annotations derived from mIF data. This analysis was conducted on six NSCLC tumors – two ISI and four ISL. We found that ISL tumors were enriched for spatial features frequently associated with immune activation including regions of necrosis and lymphoid follicular inflammation while ISI tumors were more frequently associated with perivascular inflammation, fibroblast-rich zones, and features of the tumor-stromal interface (**Fig. 5E**). To directly visualize these differences, we examined two representative tumors – one from each group. The ISL tumor (NSCLC-P6) displayed coherent spatial relationships among tumor regions, lymphoid follicles, and tumor-stroma boundaries, similar to lymphoid aggregate spatial organization in immune-responsive renal cell carcinoma (**Fig. 5F**)^77^. In contrast, the ISI tumor (NSCLC-P16) exhibited fibroblasts interlaced amongst tumor cells, an organization that has been previously linked to T cell exclusion in lung tumors (**Fig. 5G**)^78^. Thus, our results demonstrated that the ISL and ISI designations encompass known structural hallmarks of ICB responsiveness in patients.

While the ISL and ISI categories were defined using pan-tumor information, the clinical association with response was derived from our NSCLC cohort. We therefore sought to identify whether a subset of pan-tumor immune lability genes might be specifically responsible for the clinical relevance of ISL and ISI distinctions in NSCLC. To do this, we identified the genes with differential SLAB between the pan-tumor ISL/ISI cohorts, and then performed a differential SLAB analysis of these genes in NSCLC ISL versus ISI tumors. Of these, the top genes distinguishing ISL from ISI in NSCLC included STARD7 and HINT1, genes that have been implicated in TME immunomodulation across diverse cancers that include lung adenocarcinomas and lung squamous cell carcinomas (**Fig. 5H**, left)^79,80^. ORA pathway enrichment of the genes distinguishing ISL from ISI in NSCLC was notable for 1) the well-studied KEAP1/NFE2L2-axis of tumor immunosuppression in lung tumors and 2) metabolism of linoleic acid, a member of the polyunsaturated fatty acid class that has been shown to increase ICB sensitivity in NSCLC tumors (**Fig. 5H**, right)^81,82^. On the other hand, pathways that distinguished ISL from ISI in the pan-tumor cohort but not our NSCLC cohort were notable for insulin-like growth factor (IGF) signaling. Although IGF1R signaling has been connected to tumor immunomodulation in diverse tumor types, including specifically in a study of NSCLC tumors, IGF1R alteration frequency is particularly rare in lung tumors, suggesting why it did not distinguish ISL from ISI tumors in our NSCLC cohort^83–85^.

Overall, our results illustrated that forward inference about tumor immune spatial biology using our pan-tumor cohort predicted sensitivity to ICB in an out-of-sample NSCLC cohort. Furthermore, biological characterization of ISL and ISI samples were broadly consistent with known biological factors impacting immunotherapy sensitivity.

### Spatial similarity between tumors is reproduced by a sparse panel of mIF markers

Our results illustrated that SGs are spatially, biologically, and clinically relevant organizational units of the TME. However, discovering SGs required genome-wide spatial data, a measurement that remains costly and logistically intensive in clinical settings. Therefore, a natural question is to what degree the spatial similarity between tumors defined by SGs could be measured by lower resolution measurements like mIF that might be more clinically scalable.

Common approaches for extracting information from mIF data on TMEs have included evaluating (i) the mean expression level of each marker, (ii) the relative abundance of cell types, and (iii) the relative abundance of cell neighborhoods – combinations of spatially proximal cell types at a local scale^86^. TumorSPACE is an alternate approach for identifying coherent spatial structures as it can be used for any type of data where biological information is measured and connected to spatial coordinates. By applying TumorSPACE to mIF data, we defined SGs based on marker abundance and computed marker SLAB scores that captured how each marker spatially varied across SGs within a tumor (**Fig. 6A**, left) (Methods). This enabled us to ask which representation of mIF data most faithfully recapitulated the spatial similarity between samples defined by genome-wide SLAB profiles.

**Figure 6.**
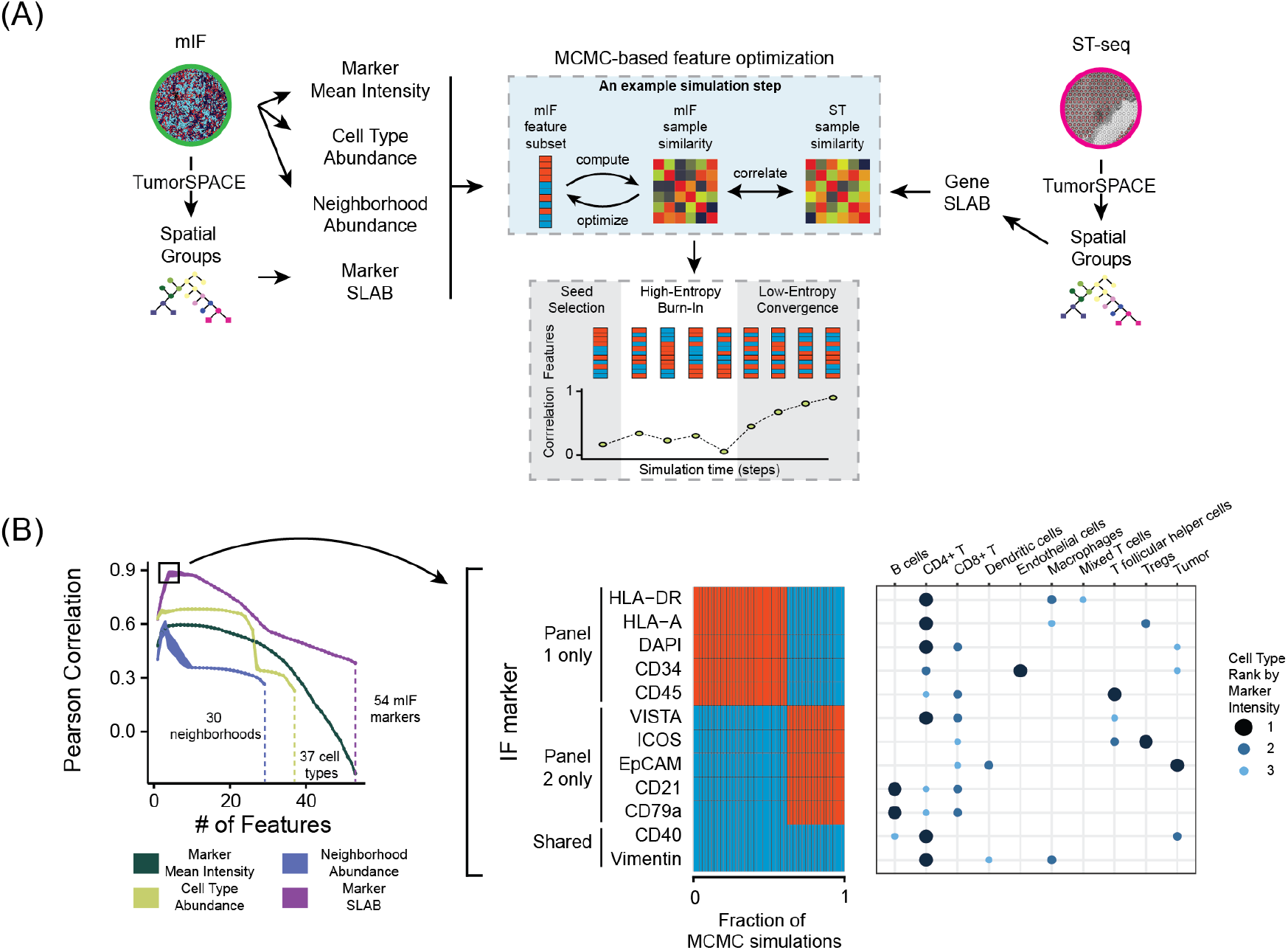
Sparse sets of IF markers capture spatial similarity between tumors. **(A)** Workflow for Markov Chain Monte Carlo (MCMC) optimization process. Left side shows that mIF data is used to directly create three different representations of tumor spatial biology: marker mean intensity, cell type abundance, neighborhood abundance. TumorSPACE performed on mIF data defines SLAB scores for each marker as a fourth representation: marker SLAB. Each of these four representations is evaluated for the capacity to capture the spatial similarity between NSCLC tumors as determined by genome-wide SLAB scores (right) through a MCMC computer simulation (middle). The essence of the simulation is an iterative search (blue box) where, in each step, mIF-based representations of spatial similarity are evaluated against spatial similarity determined by genome-wide SLAB using Pearson correlation (y-axis in grey box). Across simulation time (grey box), there is an initialization step, a high-entropy burn-in phase, and a low-entropy phase that ends when the Pearson correlation is no longer significantly different between simulation steps. **(B)** (Left) Pearson correlation (y-axis) between spatial similarity determined by various mIF representations (color key) and spatial similarity determined by genome-wide SLAB versus number of features considered (x-axis). (Right) The two top-performing panels used marker SLAB with Pearson correlation of ∼ 0.9. Each panel consists of five distinct markers and two shared markers (CD40 and vimentin) Heatmap indicates marker presence (blue) or absence (red) amongst 50 independent MCMC simulations (x-axis). For each marker, the right table indicates the top three cell types (x-axis) ranked by intensity of that marker in our mIF data (color key).

We leveraged the subset of six NSCLC tumors for which we had aligned ST-seq data and mIF data (Methods). Each of the four mIF data representations – marker mean intensity, cell type abundance, neighborhood abundance, and marker SLAB – contains a discrete set of unique features. We did not want to assume that all features were relevant for tumor spatial biology. Given the many-to-many mapping between such features and biology in the TME (e.g. tumors being enriched for both E-cadherin and EpCAM, while EpCAM is enriched on both tumor cells and dendritic cells), we hypothesized that many different or overlapping ‘feature subsets’ could recapitulate tumor biology to a certain degree. We therefore sought a data-driven approach for identifying the optimal subset of features. We created a Markov Chain Monte Carlo (MCMC) simulation framework that probabilistically sampled combinations of features across each of the four mIF-based representations. The procedure begins by selecting a random subset of features referred to as ‘the seed’. From there, the MCMC algorithm enters a phase of broad exploratory sampling across the space of all features, termed ‘high-entropy burn-in’. This phase allows the system to escape locally optimal feature combinations and instead survey a wide range of possibilities. Once promising regions of the search space are identified, the algorithm transitions to a more focused ‘low-entropy convergence’ phase that fine-tunes the selection of features by exploring nearby combinations. At each step in the MCMC algorithm, the selected feature subset is evaluated against a predefined target function: the Pearson correlation between the tumor-by-tumor similarity matrix computed using the mIF feature subset and the spatial similarity matrix computed from genome-wide SLAB scores. The simulation terminates once further changes to the feature subset result in minimal changes to this correlation (**Fig. 6A**) (Methods).

The MCMC simulation revealed that marker SLAB outperformed all other mIF representations. Notably, the correlation using marker SLAB peaked with seven features, after which performance declined. In contrast to marker SLAB, the three other representations of mIF data performed worse, displaying lower overall correlation as well as performance degradation with increasing feature count (**Fig. 6B**, left). We next identified which specific markers underpinned the ability of marker SLAB to recapitulate genome-wide SLAB similarities between the NSCLC tumors. We found two marker panels composed of seven markers each (five distinct markers and two overlapping markers: CD40 and vimentin) that were the only two solutions across a collection of 50 independent MCMC simulations (**Fig. 6B**, right). We mapped the expression of each marker across various cell types and found that both panels were enriched for markers specific to similar immune and stromal populations including CD4+ T cells, CD8+ T cells, macrophages, regulatory T cells (Tregs), and tumor cells (**Fig. 6B**, right). This result indicated that while the two marker panels were compositionally distinct, they encoded similar distributions of cell types.

To learn whether these two marker SLAB panels depended on capturing collective information across their constituent markers, we alternatively selected marker SLAB panels using a generalized linear model (GLM) regression framework. Since GLMs assess the contribution of each individual marker in isolation, we selected the top seven independently performing markers to constitute our GLM-selected marker panel (**Fig. S14A**). The GLM-selected marker panel did not fully overlap with either MCMC marker panel. Moreover, several markers in the MCMC marker panels had relatively modest GLM coefficients. While the MCMC marker panels recapitulated genome-wide SLAB-based spatial similarity between tumors (r = 0.92), the GLM-selected marker panel yielded lower concordance (r = 0.59) (**Fig. S14B,C**). These results signified that finding the optimal marker SLAB panel required considering collective information between markers and not only independent effects.

Together, these findings demonstrated that the spatial architecture defined by genome-wide SLAB profiles can be approximated using a sparse set of protein markers from mIF data. Additionally, we showed that in order to recover the necessary information from mIF data, it was important to identify mIF-based SGs and compute SLAB from markers directly.

## Discussion

Our results establish Spatial Groups (SGs) as TME spatial domains that are directly inferred from data and represent meaningful biological and histological structures. By examining SGs across diverse tumor types, we uncovered a broadly conserved principle of TME spatial biology: the organization of SGs is a nested hierarchy reflecting context-informed biological heterogeneity. Comparing TME architecture defined by SGs across hundreds of tumors revealed that the primary axis of spatial variation was immune biology. This observation resulted in the creation of a spatially aware biomarker stratifying response to immune checkpoint blockade (ICB) therapy in a retrospective cohort of NSCLC patients.

Several limitations should be considered when interpreting our results. First, while we defined nine Spatial Classes from our pan-tumor cohort, we acknowledge that our cohort does not represent all cancer types and patient populations. This motivates the question: are there additional Spatial Classes we have not properly accounted for? Our data showed that some tumor types comprise their own Spatial Class and that some Spatial Classes were only discoverable when studying a large group of tumors. These observations illustrate that while additional sampling of tumors is currently needed, it will be important for the field to consider the point at which new Spatial Classes are unlikely to be discovered with additional data. Second, our analysis intersected only two types of spatial information: RNA abundance through transcriptomics and protein abundance through mIF. Future studies could incorporate other descriptions of spatial biology (e.g. metabolomics) to assess how these complementary data provide additional context to TME spatial biology. Finally, the NSCLC cohort we evaluated was modest in size. While this sampling was sufficient to prove the concept of deriving clinically useful spatial biomarkers, our results motivate pursuing a multi-site prospective trial to test whether SG-based classifications are better than the current standard (PD-L1 IHC) as a predictive clinical biomarker of frontline ICB response.

The conserved organization of SGs into nested hierarchies that span multiple scales of biological information enables a clear direction for the field of high-resolution tumor spatial biology. Currently, our capacity to collect such data far exceeds the rate at which human expert analysis could possibly analyze these data. This necessitates establishing artificial intelligence approaches that are aligned with principles of tumor spatial biology. Our results demonstrate the integration of biology across spatial scales in the TME, thus suggesting that a natural choice of model to align with this pattern of biological organization would be a neural network. The default structure of such models is to allow for all-to-all connections between neurons of adjacent layers, implying a complete lack of constraint. However, recent work has demonstrated that incorporating constraint into models of statistical learning is critical and has led to the widespread adoption of deep learning models in many fields^87–89^. Could the hierarchy of SGs be a reasonable set of constraints that enable the success of such models in the field of tumor spatial biology? Further work could explore this idea by explicitly modeling SG connections as connected neurons in a manner inspired by ‘biologically-aware’ neural networks^90^.

In this work, we have developed a unified coordinate system for comparing tumors based on their spatial organization irrespective of tumor origin and data source. Additionally, using our NSCLC cohort as proof-of-concept, we have shown that new tumors profiled by other groups could be embedded into this space to learn about the spatial biology of those tumors as informed by our entire pan-tumor cohort and subsequently develop clinically actionable spatial biomarkers. These results suggest that as more datasets are projected into the pan-tumor space, particularly those with linked clinical annotations, the collective reference can grow in power and sharpen our ability to interpret tumors in clinically meaningful ways. As any single laboratory likely does not have the resources to perform ST-seq on clinically annotated samples at scale, we imagine that the unified coordinate system we have defined here could ultimately serve immense utility as a publicly available resource. In service of this goal, our approach enables a decentralized framework aiding in the sharing of protected clinical data across institutions that is often restricted by regulatory, security, and governance barriers. With our framework, each institution can map its tumors onto a common coordinate system locally and subsequently report and exchange only the derived coordinates and variable spatial features. This allows multi-center comparison, validation, and discovery without moving or exposing patient-level data while simultaneously leveraging ever-growing collective knowledge to enrich our understanding of TME spatial biology and clinical application.

TME spatial biology encompasses a wide range of biophysical processes that impact each other in ways we do not yet understand. As the toolbox of spatial-omics technologies continues to expand, it will become increasingly important to relate biological learnings from these technologies that directly correspond with each other. Our results have demonstrated a complex but definable mapping between a high-dimensional spatial modality (Visium ST-seq) and a low-dimensional spatial modality (CODEX mIF) that made no assumptions about the biological content of each dataset. These results open the door for SG-guided clinical trials whereby genome-wide spatial biology is captured by sparse diagnostic panels. They also suggest that SGs occupy a central place in TME spatial biology that spans RNA and protein abundance and that may span additional biological axes governing the spatial organization of the TME. Our findings underscore the importance of understanding the origins of SGs and the diverse mechanisms by which they are maintained and evolve.

## Methods

### Computational method details

#### Download and harmonization of ST-seq data

Previously deposited ST-seq datasets (**Supplementary Table 1**) were downloaded for integration from GEO (https://www.ncbi.nlm.nih.gov/geo/) into the pan-tumor ST-seq database as long as they had the following SpaceRanger outputs available: 1) a spot-by-UMI gene count matrix, 2) a spot-by-pixel location matrix, 3) a scalefactors_json.json file containing ‘spot_diameter_fullres’, and 4) the associated H&E image stored either as “tissue_hires_image.png”, “tissue_lowres_image.png”, or a full-resolution image. For analyses including physical distance rather than pixel distance, pixel distance was converted to physical distance by computing a 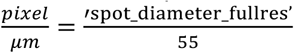 scaling factor that compares spot diameter in pixels to the known spot diameter of 55 *μ m*.

#### SpaceRanger processing of ST-seq data

For internally generated ST-seq datasets, reads were aligned and mapped to the hg38 (GRCh38) human genome reference using the SpaceRanger v2.0.0 count pipeline (**Supplementary Table 8**). This pipeline generates a raw unique molecular identifier (UMI) gene count matrix in which each row consists of a spot that has X/Y coordinates in pixels that correspond to the aligned H&E image. The SpaceRanger algorithm also identifies spots within or outside of detectable tissue, and for all subsequent analyses only spots within tissue were used.

#### Pathologist annotation of ST-seq H&E data

Whole-slide H&E images were collected as part of an ST-seq experiment (see ‘*10X Visium CytAssist spatial…’*) or as part of a publicly deposited ST-seq dataset (see ‘*Download and harmonization of ST-seq data’*). Image data were loaded into QuPath (version 0.4.4) for visual inspection and manual annotation by a board-certified pathologist. An initial review of all slides revealed a natural hierarchy of histologic features, which was used to develop a structured annotation scheme (**Fig. S2A, Supplementary Table 9**). Feature categories were defined based on the following criteria: (1) biological relevance, (2) generalizability across tumor types, (3) consistency with a hierarchical annotation scheme, and (4) non-redundancy.

All tissue-containing regions were manually annotated in their entirety. Regions of interest (ROIs) were labeled according to the predefined hierarchical scheme to ensure consistency across the dataset. Ambiguous regions were reviewed in consultation with a second pathologist. In addition, a random subset of 10% slides was re-reviewed by the primary pathologist several months after the initial annotation, with blinded access to prior labels, which confirmed intra-observer reproducibility. Label annotations were identified for ST-seq spot locations by 1) importing ST-seq spot centroid locations into QuPath and 2) identifying the annotated region containing any given spot. These analyses were performed using custom Groovy scripts that have been made available within the figure repository for this paper (see ‘*Code availability’*).

#### TumorSPACE: models and associated analysis

The sub-sections within this section will introduce several variables. Below is a table of all variable definitions.

| <u>Variable</u> | <u>Definition</u> | <u>Section Where First Referenced</u> |
| --- | --- | --- |
| *M* | SpaceRanger UMI gene count matrix | *Creating a latent space* |
| $m$ | Number of ST-seq spots in $M$ | *Creating a latent space* |
| $n$ | Number of genes in $M$ | *Creating a latent space* |
| $U$ | SVD left singular matrix | *Creating a latent space* |
| $\Sigma$ | SVD singular value matrix | *Creating a latent space* |
| $V^T$ | Transpose of SVD right singular matrix | *Creating a latent space* |
| $D$ | Spectral Distance matrix | *Creating a latent space* |
| $p$ | PC depth hyperparameter | *Creating a latent space* |
| $T$ | TumorSPACE tree model | *Creating a latent space* |
| $G$ | The set of tree internal nodes | *Creating a latent space* |
| $g$ | A single tree internal node | *Creating a latent space* |
| $M_B$ | Bootstrapped gene count matrix | *Bootstrapping the latent space...* |
| $X$ | Statistical random variable | *Bootstrapping the latent space...* |
| $N$ | Normal Distribution | *Bootstrapping the latent space...* |
| $\mu$ | Mean | *Bootstrapping the latent space...* |
| $\sigma$ | Standard deviation | *Bootstrapping the latent space...* |
| $T_B$ | Bootstrapped TumorSPACE tree | *Bootstrapping the latent space...* |
| $b$ | Node TBE support | *Bootstrapping the latent space...* |
| $k$ | Node spatial dispersion | *Calculating physical spatial dispersion...* |
| $K$ | Ripley's reduced second moment function with border correction | *Calculating physical spatial dispersion...* |
| $g_s$ | The set of spots in node $g$ | *Calculating physical spatial dispersion...* |
| $len_x$ | The x-axis range of values for a set of spots | *Calculating physical spatial dispersion...* |
| $len_y$ | The y-axis range of values for a set of spots | *Calculating physical spatial dispersion...* |
| $\lambda$ | Ripley intensity normalization factor | *Calculating physical spatial dispersion...* |
| $r_{max}$ | Ripley maximum spot distance | *Calculating physical spatial dispersion...* |
| $R$ | The set of spot distances used to compute spatial dispersion | *Calculating physical spatial dispersion...* |
| $r$ | A particular spot distance for computing spatial dispersion | *Calculating physical spatial dispersion...* |
| $\delta$ | Spot pairwise physical distance | *Calculating physical spatial dispersion...* |
| $\Delta$ | Matrix of pairwise spot-spot physical distances | *Calculating physical spatial dispersion...* |
| $X_{spot}$ | X axis physical location of a spot | *Calculating physical spatial dispersion...* |
| $Y_{spot}$ | Y axis physical location of a spot | *Calculating physical spatial dispersion...* |
| $\kappa$ | The number of spot KNN matches | *Calculating physical spatial dispersion...* |
| $P$ | The set of $p$ being tested | *Calculating physical spatial dispersion...* |
| $n_p$ | The length of set $P$ | *Calculating physical spatial dispersion...* |
| $Uniform([a,b])$ | The Uniform Distribution between a and b | *Calculating physical spatial dispersion...* |
| $H$ | The set of hyperparameters for $b$, $k$, and $\kappa$ | *Calculating physical spatial dispersion...* |
| $n_H$ | The length of set $H$ | *Calculating physical spatial dispersion...* |
| $h_i$ | A given choice of hyperparameter values for $b$, $k$, and $\kappa$ | *Calculating physical spatial dispersion...* |
| $G_{filt}$ | The set of internal nodes in $G$ that meet a given set of hyperparameter bounds on $b$ and $k$ | *Calculating physical spatial dispersion...* |
| $G_{filt}'$ | The set of $G_{filt}$ plus the parent nodes of $G_{filt}$ within $T$ | *Calculating physical spatial dispersion...* |
| $T_{filt}$ | Tree $T$ filtered for internal nodes within $G_{filt}'$ | *Calculating physical spatial dispersion...* |
| $d$ | Spot pairwise spectral distance | *Prediction accuracy calculation* |
| $I$ | The set of all spots in a biopsy | *Prediction accuracy calculation* |
| $NN_{i,\kappa}$ | The set of $\kappa$ nearest neighbors to spot $i$ in latent space $T$ | *Prediction accuracy calculation* |
| $spot_{i,z}$ | A given nearest neighbor spot within $NN_{i,\kappa}$ | *Prediction accuracy calculation* |
| $\rho$ | Pearson Correlation Coefficient | *Prediction accuracy calculation* |
| $corr$ | Pearson Correlation function | *Prediction accuracy calculation* |
| $vec$ | Matrix vectorization function | *Prediction accuracy calculation* |
| $S_{spot}$ | The tuple of spot locations ($X_{spot}$, $Y_{spot}$) | *Prediction accuracy calculation* |
| $T_{filt}^{opt}$ | The optimized TumorSPACE model for a given tumor biopsy | *Prediction accuracy calculation* |
| $A_{g_i}^{path}$ | The full ancestral node path for an internal node $g_i$ in any tree $T$ | *Nested Spatial Group (NSG) depth* |
| $a_{g_i}^k$ | The $k^{th}$ ancestor of an internal node $g_i$ in any tree $T$ | *Nested Spatial Group (NSG) depth* |
| $\sigma_{node_g}$ | The spatial domain size of an internal node $g_i$ in any tree $T$ | *Nested Spatial Group (NSG) depth* |
| $\lambda_{node_g}$ | The NSG depth of an internal node $g_i$ in any tree $T$ | *Nested Spatial Group (NSG) depth* |
| $G_{DA}$ | The subset of internal nodes within tree $T$ that will be used for differential abundance computation | *SG-based differential abundance* |
| $W(a,b)$ | Wilcoxon rank-sum test between $a$ and $b$ | *SG-based differential abundance* |
| $p_{g_{DA},f}$ | Differential Abundance Probability at node $g_{DA}$ for process $f$ | *SG-based differential abundance* |
| $q_{g_{DA},f}$ | Differential Abundance Probability at node $g_{DA}$ for process $f$, empirically bootstrapped and multiple hypothesis adjusted | *SG-based differential abundance* |
| $DA$ | The set of differentially abundant spots for process $i$ at node $j$ in a given tumor biopsy | *SG-based differential abundance* |
| $OR_{i,j}$ | The odds ratio of independence between process $i$ in an NSG and process $j$ in a parent SG | *Contextual dependence of processes...* |
| $SLAB$ | The SLAB Score for process $i$ in a given tumor biopsy | *SG-based spatial lability (SLAB) score* |
| $L^k$ | The matrix of SLAB scores composed of samples (rows) in groups K and nK | *Differential SLAB score analysis* |
| $MW(a,b)$ | Mann-Whitney U Test between $a$ and $b$ | *Differential SLAB score analysis* |
| $p_{K,f}$ | p-value of the MW test between samples in groups K vs nK for process $f$ | *Differential SLAB score analysis* |
| $p_{K,f}^{shuffled}$ | p-value of the MW test between samples shuffled between groups K vs nK for process $f$ | *Differential SLAB score analysis* |

##### Overview

Building a TumorSPACE model requires spatial transcriptomic data and two inputs from SpaceRanger: (1) the raw gene UMI count matrix and (2) the spot spatial coordinate matrix. Model building subsequently operates on the gene count matrix to build many models that vary in hyperparameter choice. The spot spatial coordinates are then used for selecting the optimal hyperparameter set that maximizes accurate recovery of spatial spot organization.

Four hyperparameters are tuned during this process: (1) the number of principal components (PCs) of data-variance used for creating a latent space of the transcriptional data, (2) the limit of statistical robustness for spot-spot relatedness in the latent space, (3) the spatial dispersion of the nodes in the latent space hierarchical tree model, and (4) the number of KNN matches used for spot spatial prediction. The following sections will first establish the model latent space and compute statistical robustness and spatial dispersion properties of that latent space. Subsequently, all hyperparameters will be tuned to define the optimal model for mapping transcriptional content from TME spots to TME spatial organization.

##### Creating a latent space

The first step in building a TumorSPACE model is to create a latent space representation of the gene count data that incorporates statistical bootstrapping. TumorSPACE first embeds ST-seq spots into a latent space by applying singular value decomposition (SVD) to the gene count matrix^91^:

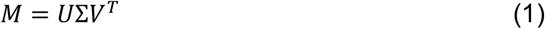

*M* is the SpaceRanger gene count matrix (*m* spots as rows, *n* genes as columns), *U* is the left singular matrix, *Σ* is the singular value matrix, and *V* is the right singular matrix. *U* is defined by cell spots (rows) and left singular vectors (columns), where each entry is the projection of a cell spot onto a left singular vector. *Σ* is a diagonal matrix where entries are singular values. *V*^*T*^ is defined by genes (rows) and right singular vectors (columns) where each entry is the projection of a gene onto a right singular vector.

First, from (1), a metric termed ‘spectral distance’ (*D*) between all spots is calculated. This metric was previously developed by our laboratory in the context of analyzing phylogenetic bacterial proteome content^28^. As implemented for spatial data in this manuscript, performing SVD on the gene count matrix determines the extent to which each cell spot projects onto each left singular vector. Therefore, a distance considering the transcriptomes of two spots can be computed by measuring the difference in the projections of two spots onto a left singular vector. Note, this definition of distance does not consider any information about spatial spot distribution.

Next, groups of left singular vectors are combined to create ‘spectral groups’. These groups are defined based on the eigenvalues associated with each left singular vector: left singular vectors with similar eigenvalues are grouped together:

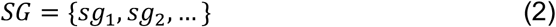

where *SG* is the total set of spectral groups, *sg*_1_ is first set of columns extracted from *U, sg*_2_ is the second set of columns extracted from *U*, and so on. The concept of spectral groups was also previously developed by our laboratory^39^. Defining *sg*_1_ and *sg*_*i*+1_ is done by identifying larger than expected decreases in singular values between consecutive left singular vectors. To compute spectral groups, a vector of differences between consecutive singular values is computed for all left singular vectors. We use the upper and lower quartiles of this distribution in combination with a scaling parameter alpha to define the ‘expected difference’ bounds between singular values. Any difference in singular values outside of these bounds deviates from expectation and therefore defines a spectral group (see associated GitHub code for specification of parameters). The spectral distance for a pair of spots within a spectral group is then computed as the Euclidean distance between spot projections onto left singular vectors comprising the spectral group weighted by the eigenvalue associated with each left singular vector. The summation of these distances across all spectral groups is the spectral distance, *d*_*i,j*_, between spots *i* and *j*.

After computing *d*_*i,j*_ for all spots, the resulting construct is a spectral distance matrix *D* comprised of *m* rows and *m* columns where *m* is the number of spots in the original gene count matrix and each entry in *D* is the spectral distance between two spots. *D* is then used as input for hierarchical clustering with complete linkage, resulting in a tree *T* that relates all spots in a tumor sample to each other. *T* has *m* leaves and (*m-1*) internal vertices (nodes). The leaves are the ST-seq spots and the nodes *g* ∈ *G* represent *G* hierarchically ordered groupings of these spots. The resulting network is the TumorSPACE latent space of the original gene count matrix.

The number of spectral groups is dependent on how many of the total left singular vectors are considered. An increasing number of left singular vectors being included corresponds directly to the inclusion of deeper principal components when computing the latent space. For TumorSPACE models, the depth of principal components, ‘*p’*, is a hyperparameter that is tuned for embedding the gene count matrix into a latent space.

##### Bootstrapping the latent space to evaluate statistical robustness

TumorSPACE does not assume that each node *g* arises from biological signal. Instead, TumorSPACE bootstraps *T* using the Booster package’s implementation of transfer bootstrap expectation (TBE), the probability that node *g* appears in an empirically bootstrapped tree (default settings used for Booster)^92^. For generating empirically bootstrapped trees, we applied Gaussian multiplicative noise injection to the initial gene count matrix *M* to create a “bootstrapped” gene count matrix *M*_*B*_.

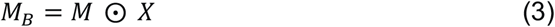

such that ⊙ indicates element-wise multiplication by a normally distributed random variable *X* ∼ *N*(*μ, σ*^2^) with μ = 1 and *σ* = 0.2. This matrix was then used as an input to (1) and a tree was created following the steps outlined in ‘*Creating a latent space*’ to generate a bootstrapped tree *T*_*B*_. Bootstrapping was done 10 times for a given dataset, followed by input of the original tree *T* and the bootstrapped trees *T*_*B*_ into Booster for TBE computation. This results in a labeling of the original tree *T*’s set of nodes *G* with TBE support values *b*_*G*_ such that *b*_*G*_ ∈ [0,1].

##### Calculating physical spatial dispersion in latent space

The final property of *T* that is computed is the spatial dispersion *k* for each node comprising *T*. Spatial dispersion is estimated for each node using Ripley’s reduced second moment function *K(r)* with border correction^93,94^. Let *g*_*s*_ be the set of ST-seq spots within node *g* in *T*. The window of physical tumor space is defined by the spot spatial coordinate matrix such that *len*_*x*_ indicates the x-axis window length and *len*_*y*_ indicates the y-axis window length. We then compute, a normalization factor for spot intensity within a spatial region, and *r*_*max*_, a factor that incorporates lambda to determine the maximum spatial distance being assessed.

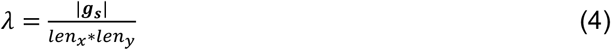

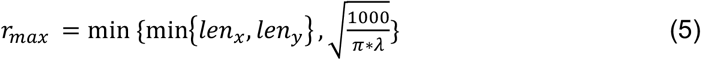

where *min* denotes the minimum between a set of values. Let *R* be the set of spot spatial distances that will be assessed, such that

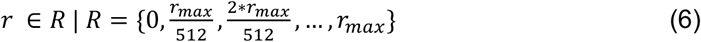

We define the physical distance *δ* between any two spots as

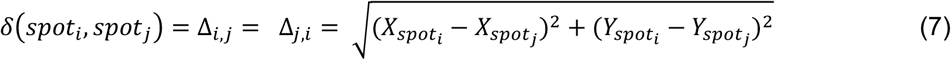

where (X_spot_,Y_spot_) denote the physical space coordinates for a given spot. Spatial dispersion *K(r)* with border correction is then computed for all spots *g*_*s,i*_, ∈ *g*_*s*_ as

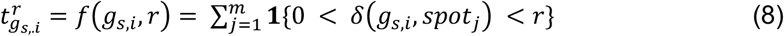

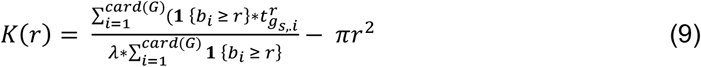

where *t*(*g*_*s,i*_, *r*) is the number of spots within distance *r* of a given *g*_*s,i*_ and *b*_*i*_ is the distance from spot *g*_*s,i*_ to the window boundary. The general notation *card(S)* indicates the number of elements in a set S, and the general notation ***1****{f(x)}* signifies a value of 1 when f(x) is true and a value of 0 when *f(x)* is false. Finally, spatial dispersion *k* is computed by summing the absolute value of *K(r)* over *r* ∈ *R* as follows.

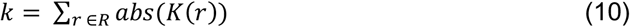

This calculation labels all nodes *G* in tree *T* with spatial dispersion values *k*_*G*_ such that *k*_*G*_ ∈_ℝ_ [0, ∞].

##### Hyperparameter optimization to create a TumorSPACE map

TumorSPACE model optimization involves selecting the values of four hyperparameters that maximize model prediction accuracy (described in ‘*Prediction Accuracy Calculation*’) for a given dataset. These hyperparameters tune three properties of tree *T* – principal component depth *p* (from ‘*Creating a latent space*’), node TBE support *b* (from ‘*Bootstrapping the latent space to evaluate statistical robustness*’), node spatial dispersion *k* (from ‘*Calculating physical spatial dispersion in latent space*’) – as well as one property of accuracy computation, the number of spot KNN matches κ. We perform hyperparameter tuning as a nested grid search by tuning *p* as an outer layer and then optimizing [*b, k, κ*] for a given value of *p*.

First, a set of PC depth values (*n*_*p*_ where default is set to 10) is randomly selected to create a set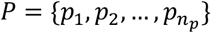. The PCs termed *p*_*i*_ are chosen on a logarithmic interval between a minimum and maximum PC depth, which is the rank of the gene count matrix *M*. Next, a matrix of three hyperparameter values, 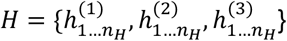, are created where the vectors *h*^(1)^, *h*^(2)^, and *h*^(3)^ are independently sampled from distributions as follows.

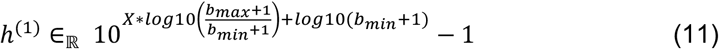

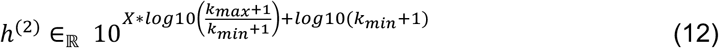

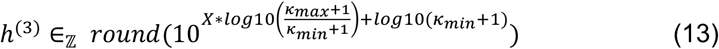

In (*11-13*), X is a random variable drawn from *Uniform*([0,1]). Default values for hyperparameter bounds are *b*_*min*_ = 0, *b*_*max*_ = 0.5, *k*_*min*_ = 0, *k*_*max*_ = 1 κ_*min*_ = 5, κ_*max*_ = 300. A minimum of *n*_*H*_ = 100 sets of {*h*^(1)^, *h*^(2)^, *h*^(3)^} are initially sampled, after which additional sets are sampled until prediction accuracy optimization has converged. Prediction accuracy convergence is reached when the difference in prediction accuracy (defined below in ‘*Prediction Accuracy Calculation’*) for the top 2 scoring hyperparameter sets is less than 0.05. For a given hyperparameter set *h*_*i*_, the TumorSPACE tree *T* is filtered for the set of nodes *G*_*filt*_ such that each node in *G*_*filt*_ satisfies

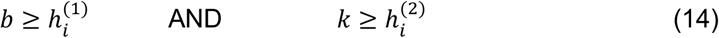

The final filtered tree, *T*_*filt*_, comprises the set of nodes *G*_*filt*_*’*, which consists of *G*_*filt*_ as well as the complete set of parent nodes from which *G*_*filt*_ descend even if those parent nodes do not meet the criteria in *(14)*, along with all ST-seq spots. We found that optimal hyperparameter values varied widely amongst datasets, underscoring the importance of fitting these parameters for each individual dataset (**Fig. S15**).

##### Prediction accuracy calculation

To identify the TumorSPACE model properties that were optimized for predicting spot spatial locations from transcriptomic data, we masked the physical location of each ST-seq spot and identified its *k* nearest neighbors in the TumorSPACE latent space by minimizing spectral distance.

For any masked spot *i* amongst all spots *I*, we can define its κ nearest neighbors *NN*_*i,k*_ as

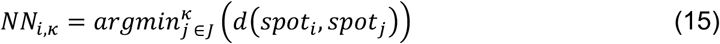

where *i* ∈ *I*, κ ∈ *h*^(3)^ as defined in (*13*), *J* is the set of all spots other than spot *i*, and *argmin*^*κ*^ selects the set of κ spots with the smallest spectral distance relative to spot *i*. To prevent overfitting, we identified for each *spot*_*i,z*_ ∈ *NN*_*i,k*_ a randomly chosen 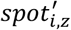 that belongs to the internal node *g*^*i,z*^ within *T*_*filt*_ immediately ancestral to *spot*_*i,z*_.

We then estimated the location of masked *spot i* based on the x and y locations of the corresponding 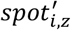 spots.

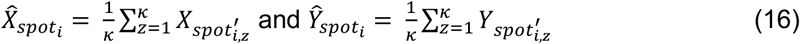

Finally, we computed the Pearson Correlation *ρ* between the vectorized matrix *Δ*_*actual*_ of pairwise actual spot-spot physical distances and the vectorized matrix *Δ*_*predicted*_ of pairwise predicted spot-spot physical distances.

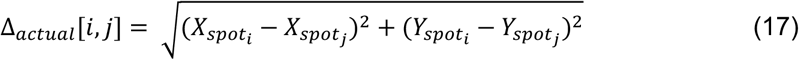

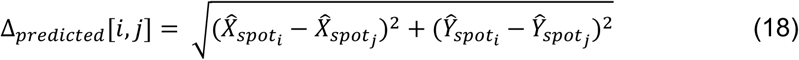

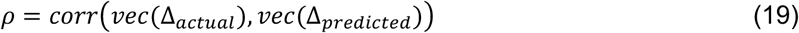

where *vec()* indicates matrix vectorization to a single column and *corr()* indicates Pearson Correlation. To compute a null distribution for *ρ* using empirical bootstrapping of actual versus predicted spot locations in a given dataset, we shuffled the vector 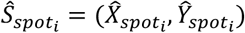 without replacement and then re-computed *(17-18)* using this shuffled vector 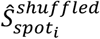 of predicted spot locations.

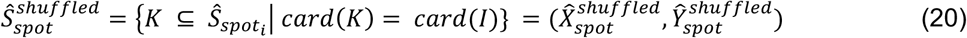

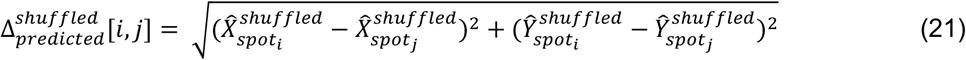

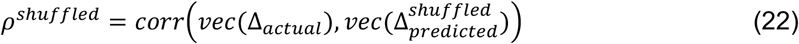

For **Fig. 1C**, *ρ*^*shuf fled*^ is computed for 100 shuffles and the maximum *ρ* is taken as the ‘null’ prediction value. The null distribution is plotted in the grey distribution in **Fig. 1C**.

Finally, the optimal TumorSPACE model 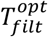 is found that maximizes *ρ* across hyperparameter sets *P* and *H*.

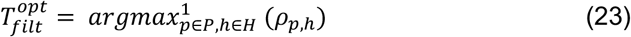

##### TumorSPACE model outputs

For a given input tumor ST-seq dataset, the output from TumorSPACE includes: (1) the TumorSPACE model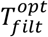, (2) the Pearson Correlation estimate *ρ*, and (3) the set of predicted spot locations 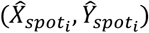 for all ST-seq spots. The final set of internal nodes within 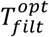 are termed Spatial Groups (SGs).

##### Nested Spatial Group (NSG) depth

We computed NSG depth as a measurable quantity that describes how a given NSG relates to the other parts within a TumorSPACE model. As such, we define ‘NSG depth’ as a property of all NSGs within a TumorSPACE model.

To first define NSG depth, we compute the complete ancestral node path for any internal node *g*_*i*_ within 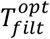 as

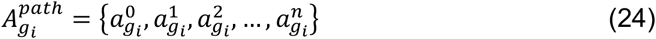

such that

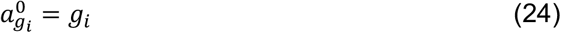

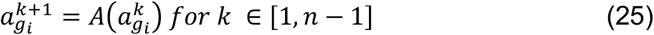

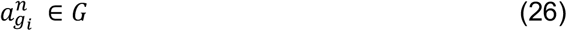

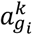 indicates the *k*^*th*^ ancestral node of node *g*_*i*_, *A(node)* denotes the immediate ancestral node of a given node in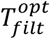, and *G* is the set of internal nodes in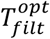. By definition, 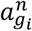 will be the root node of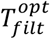. We next define the spatial domain size 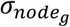 for a given node *g* as the mean spot-spot physical distance between all spots within *g*.

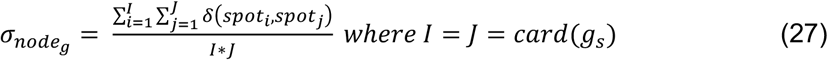

Finally, we identify the subset of nodes 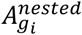 within 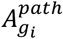 that satisfy the condition whereby the *(k+1)*^*th*^ node is equal to or larger in spatial domain size than the *k*^*th*^ node in that path.

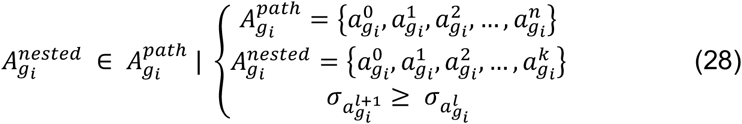

where *k* ≤ *n* and 0 ≤ *l* ≤ *k*. The NSG depth, 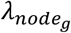, for a given internal node *g*_*i*_ is defined to be the number of ancestral generations that satisfy this condition of spatial domain nesting.

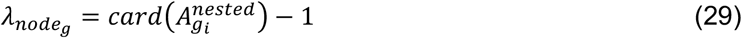

##### SG-based differential abundance

Differential abundance calculation requires two inputs: (1) an optimized TumorSPACE model 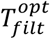 and (2) a spot-by-feature matrix *F*. We computed differential abundance using three types of biological processes: genes, pathways, and deconvoluted cell type proportions. Computation of gene count, pathway usage, and cell type proportion matrices are described in the *‘SpaceRanger’, ‘Pathway over-representation analysis’*, and *‘SpaCET’* Methods sections, respectively. The gene count matrix is normalized by the spot-wise total UMI count.

First, we identified a subset of SGs *G*_*DA*_ ∈ *G* at which DA will be computed. We set a minimum of 10 spots that must be present in both a given SG *g*_*DA*_ ∈ *G*_*DA*_ and in its sibling node 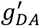(e.g. *A* and *A’* in **Fig. 3A**) for inclusion within *G*_*DA*_.

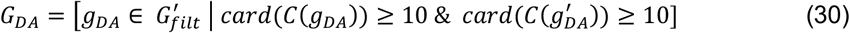

where *C(n)* indicates the row indices within matrix *F* of the spots descending from SG *g*_*DA*_. Subsequently, for each node *g*_*DA*_ and process *f*, the spot-wise process values between *g*_*DA*_ and 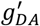 are compared using a two-sided Wilcoxon Rank Sum Test, where the test p-value is given by *W(a,b*).

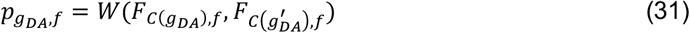

To facilitate empirical correction for multiple hypothesis testing, we perform 20 shuffles of the process values between *g*_*DA*_ and 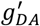, followed by computation of the Wilcoxon p-value between these shuffles. Let 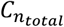 be the concatenation of spot indices *C*(*g*_*DA*_) and 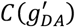.

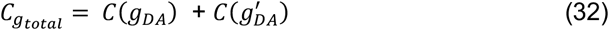

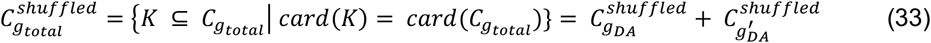

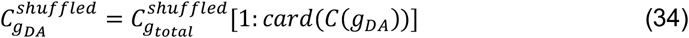

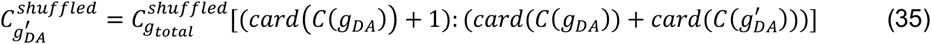

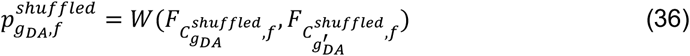

Let 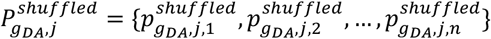 be the set of *n* DA probabilities for node *g*_*DA*_ and shuffle *j*, where *n* is the number of processes in *F*. Then, a given process is found to be differentially abundant at a given node if its unadjusted p-value, 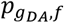, is less than the minimum of all shuffled probabilities for that node. To assign the direction of process abundance change for nodes with significant abundance changes, given that our test examines relative changes in expression between *g*_*DA*_ and in its sibling node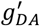, we defined the larger of the two nodes as having a “baseline expression profile” for that shared local transcriptional and spatial context. Conversely, the smaller of the two nodes was defined as having either increased or decreased abundance relative to the larger node.

##### Contextual dependence of processes based on architecture of SGs

To determine whether differentially abundant processes within child SGs were impacted by the differentially abundant processes of their parent SGs, we computed the odds ratio test for independence as follows. Let *f*_*i*_ ∈ *F* and *f*_*j*_ ∈ *F* denote two biological processes drawn from the set of all pathways and cell types identified (see *‘Pathway over-representation analysis’* and *‘SpaCET’* sections). Across all TumorSPACE models, we identified the set of child-parent SG pairs – denoted by (*N*_*i*_, *P*_*j*_) – such that 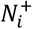 and 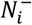 indicate the subset of child SGs where process *i* was increased or decreased in abundance, respectively, and 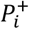 and 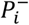 indicate the subset of parent SGs where process *j* was increased or decreased in abundance, respectively. Then, the odds ratio of independence *OR*_*i,j*_ was defined as,

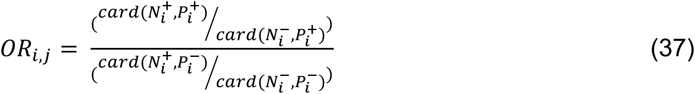

Standard definitions were used for calculation of odds ratio standard error and p-values^95^.

##### SG-based spatial lability (SLAB) score

Given a single TumorSPACE model 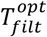 and a process *f*_*i*_ for which differential abundance has been computed in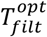, we define the SLAB score as follows. Let 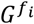 be the set of SGs in 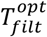 in which the process *f*_*i*_ is differentially abundant (*q* < 0.05). For each node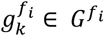, this means that process *f*_*i*_ is differentially abundant between 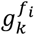 and its sister node 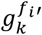 in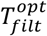. First, we identify which of the nodes, either 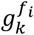 or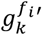, contains the fewer number of spots. This node is defined as the node with either increased or decreased abundance of process *f*_*i*_, while the node with the greater number of spots is considered to be the ‘baseline’ abundance state for process *f*_*i*_ in that subset of the tumor biopsy. 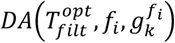 describes the set of spots with differential abundance in process *f*_*i*_ for TumorSPACE model 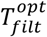 at node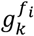:

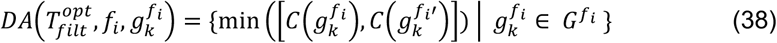

Next, we compute the union of those differentially abundant spots and compute the fraction that these spots constitute compared to the total set of spots *I* in the biopsy as a whole.

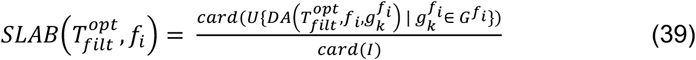

##### SLAB score correlation with bulk expression

For correlation of genome-wide SLAB scores with bulk gene expression, as in **Fig. 3B** and **Fig. S9A**, we did the following. First, we identified the set of all dataset-gene pairs for which the gene had greater than 0 UMIs detected per spot and a non-zero SLAB score in that dataset. Next, to enable computing correlation statistics, we identified genes with greater than 10 dataset entries in the filtered dataset-gene pair list. For these genes, we computed the Pearson Correlation estimate and p-value between SLAB score and mean spot UMI count across datasets. Correction for multiple hypothesis testing was done using the Benjamini-Hochberg method with a corrected q-value threshold of 0.05^96^.

##### Spatial lability pan-tumor classification

Given the set of genome-wide SLAB scores that were computed for all datasets within the pan-tumor ST-seq database (see ‘*Download and harmonization of ST-seq data*’ and ‘*SpaceRanger processing of ST-seq data*’), we aligned these score vectors into a matrix 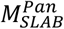 such that each dataset was a row and each gene was a column. For any instances where a gene had mapped reads in one dataset but not another – thus resulting in blank cells in this matrix – the score within this matrix was set to zero. Next, Euclidean distance was computed between each pair of rows, resulting in a distance matrix that compared all datasets to each other. Finally, the Unweighted Pair Group Method with Arithmetic mean (UPGMA) algorithm was used for constructing a hierarchical tree relating datasets to each other (**Fig. S9C**)^97^.

Spatial Classes were identified from this hierarchical tree by filtering the tree branchpoints in two steps. First, only branchpoints with at least five datasets in each child branch were retained. Second, empirical bootstrapping using transfer bootstrap estimation (TBE) was applied to identify statistically robust branchpoints^98^. For this, the SLAB matrix was re-sampled 100 times so that columns (genes) were randomly selected with replacement. From these re-sampled matrices, the same hierarchical tree algorithm was applied, resulting in 100 bootstrapped trees. The original tree was then compared to these re-sampled trees to infer TBE support values ranging from 0 (no reproducibility) to 1 (perfectly reproducible) for each branchpoint in the tree. Branchpoints with TBE >= 0.95 were retained at this step. The resulting terminal leaves of these branches were defined as ‘Spatial Classes’.

In order to simulate defining a pan-tumor classification of tumors based on SLAB scores using variably sized training cohorts consisting of randomly selected tumors (see **Fig. S10B**), we did the following. Our pan-tumor cohort consisted of 202 unique tumors amongst the 246 ST-seq datasets. We simulated 100 random orderings in which these 202 tumors would be added to the training data corpus one at a time. As a tumor is added to the training corpus, it is given the same Spatial Class designation as from training a pan-tumor model with all data, as in **Fig. 3E**, which are considered the ‘ground truth’ Spatial Classes for these tumors. The addition of a single tumor, along with all of its replicate datasets, constitutes one ‘epoch’ in a simulation. Following each epoch, the entire ‘testing’ set of 246 datasets is compared in their SLAB scores to each dataset within the training corpus using Euclidean distance between SLAB vectors. For any Spatial Classes represented by a non-zero number of datasets in the training corpus, the mean Euclidean distances are calculated to the testing datasets, and testing data are assigned to the Spatial Class that minimizes distance. Finally, these ‘simulated’ Spatial Class assignments are compared to the ground-truth Spatial Class assignments. Each Spatial Class is evaluated at each Epoch using an F1 statistic – the harmonic mean of precision and recall, bounded from 0 to 1.

##### Differential SLAB score analysis

At each high-confidence branchpoint identified in ‘*Spatial lability pan-tumor classification’*, we compared the datasets within the child branches for differential SLAB scores at all genes. Let the dataset groups with these branches be named *A* and *A’*.

To compare gene-level SLAB scores, we first compose the matrix *L*^*A*^ of SLAB scores where *L*^*A*^ has *a* + *a*′ rows corresponding to tumors *a* ∈ *A* and *a*′ ∈ *A*′ and *F* columns where *f* ∈ *F* constitutes the full set of genes. Next, for each gene *f*, we compare the tumors in *A* and *A’* where *C(X)* indicates the row indices within matrix *L*^*A*^ that correspond to tumors in either group. Comparison is performed using a Mann-Whitney U Test, where the test p-value is given by *MW(a,b*).

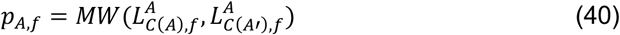

To facilitate empirical correction for multiple hypothesis testing, we perform 1000 shuffles of the SLAB counts between *A* and *A’*, followed by computation of the MW p-value between these shuffles. Let 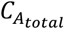 be the concatenation of row indices *C*(*A*) and *C*(*A*′).

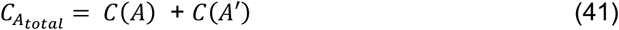

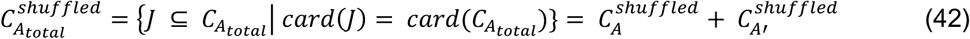

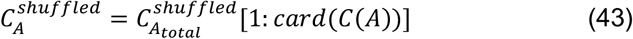

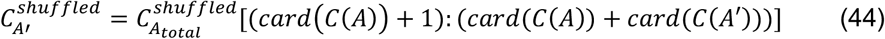

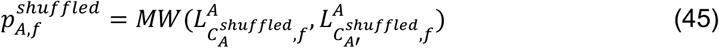

Let 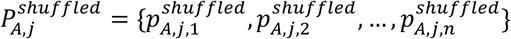 be the set of *n* probabilities for group *A* and shuffle *j*, where *n* is the number of genes in *F*. Then, a given gene is found to have a differential SLAB score between a given grouping *A* vs *A’* if its unadjusted p-value, *p*_*A,f*_, is less than the 5^th^ percentile (*q = 0*.*05*) of all shuffled probabilities.

##### Classification of NSCLC datasets by pan-tumor spatial lability

For comparison of out-of-sample NSCLC tumors to the pan-tumor Spatial Classes lability groups shown in **Fig. 3E**, we first computed SLAB scores for all genes and aligned the score vectors to match the columns (genes) of the pan-tumor SLAB score matrix 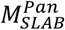. Any genes with no detectable reads for a given sample had their SLAB score set to zero. We called this new matrix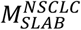. For every pair of rows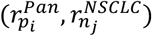, where 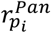 indicates the score vector for sample *p*_*i*_ ∈ *P* in the pan-tumor database and 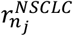 indicates the score vector for sample *n*_*j*_ ∈ *N* in the NSCLC out-of-sample dataset, we computed the Euclidean distance 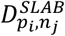 that describes the similarity between these two samples with respect to their SLAB scores. We then computed the mean of 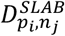 for the subset of the *P* datasets corresponding to each Spatial Class (SC1-9). Each NSCLC sample was assigned to the Spatial Class that minimized this mean distance.

##### Classification of NSCLC datasets using bulk expression and published gene sets

To determine whether classification of tumor datasets by either (1) bulk expression versus SLAB score, (2) previously published gene sets for NSCLC IO response, or (3) SpaCET-inferred cell type abundance was predictive of PFS in our NSCLC cohort, we performed the following analysis.

First, we computed aligned matrices as described in ‘*Spatial lability pan-tumor classification’* for both the pan-tumor datasets and the NSCLC datasets where matrices contained either SpaCET-inferred cell type abundance, bulk expression data or SLAB score data. For bulk expression, we computed the mean spot-wise UMI count for any given gene. Second, we filtered the aligned matrices for subset of columns (genes) described by a particular gene set or used all columns for the ‘all genes’ analysis. Third, we computed the *[16 x 246]*-dimension distance matrix between the 16 rows (samples) in the NSCLC matrix and the 246 rows (samples) in the pan-tumor matrix. Fourth, K-means clustering with K = 2 was performed row-wise on this matrix to divide the NSCLC datasets into 2 groups based on their distance vectors to the pan-tumor datasets. K-means clustering was performed 100 times for each condition using different random seeds each time. Finally, the two classes of NSCLC datasets were applied to survival analysis (described below in ‘*Survival analysis’)* to determine if they were predictive of NSCLC ICB outcomes.

#### Alignment of ST-seq and mIF data

Coordinates were aligned between CODEX mIF data and ST-seq data generated from serial tissue sections as follows. First, the DAPI channel of the CODEX data and the H&E image of the ST-seq data were identified for alignment. The alignment process employed the Scale-Invariant Feature Transform (SIFT) algorithm, which identified key points in images and matched the feature descriptors of key points to identify corresponding pairs of points from different images^99^. Based on the matched points, an affine transformation was performed to map the coordinate system of the ST-seq data to that of the mIF data. We first converted the H&E images into grayscale and adjusted their contrast using Contrast Limited Adaptive Histogram Equalization (CLAHE; implemented in OpenCV). The SIFT algorithm was then applied to detect features in both the H&E and DAPI images. Feature matching was performed based on the SIFT descriptors, with manual adjustments to the feature matching ratio depending on the similarity of the images. A transformation matrix was estimated using the Random Sample Consensus (RANSAC) algorithm, allowing for rotation, translation, and scaling transformations. The RANSAC inlier threshold was also manually adjusted to accommodate varying degrees of distortion between images. Two of the eight tissues in which alignment was attempted could not be aligned because the SIFT algorithm could not identify sufficient matching features between the images (>7 matching features).

#### SpaCET

SpaCET estimates deconvoluted cell type proportions within spots of an ST-seq experiment^100^. It requires the user to supply (1) the SpaceRanger gene count matrix as input and (2) a value for the ‘cancerType’ parameter to define the SpaCET library scRNA-seq datasets used for cell type definition. The ‘cancerType’ values chosen for each ST-seq dataset are listed in **Supplementary Table 10**. Otherwise, default parameters and commands were used as per the repository instructions (https://data2intelligence.github.io/SpaCET/articles/visium_BC.html).

#### Pathway over-representation Analysis (ORA)

For genes identified as differential between SGs (see **Figure 2**) or between Spatial Classes (see **Figure 4**), we conducted over-representation pathway analysis (ORA) for the set of Reactome pathways within the MSigDB database^101,102^ (**Supplementary Table 11**). ORA was performed using Enrichr with default parameters, which uses a Fisher exact test to compute enrichment of a gene list for a given pathway^103^. The background gene set used was the set of all genes with mapped reads in any sample. Correction for multiple hypothesis testing was implemented by using a false discovery rate threshold of < 0.1.

#### Survival analysis

For survival analysis we used the R ‘survival’ package to model progression-free survival (PFS) as a function of possible confounder variables (Treatment regimen, PD-L1 status, somatic mutation status) or classification variables (PD-L1 multi-class, PD-L1 binary, ISL/ISI, bulk expression- and SLAB score-gene sets). For confounder analysis, outcomes were modeled using Cox’s univariate or multivariate proportional hazards model. For Kaplan-Meier survival curves stratified by classification variables, survival was estimated using the Kaplan-Meier method and reported p-values were calculated using the log rank statistical test. For censored data labeling, 1 indicates that PFS was observed while 0 indicates the patient was censored for PFS.

#### Pathologist annotation of CODEX mIF data

##### Nuclear segmentation and cell boundary definition

Following image acquisition and pre-processing (see ‘*Experimental method details: CODEX multiplexed immunofluorescence’*), we applied the neural network-based cell segmentation tool, DeepCell, on the DAPI channel for nuclei identification^104^. Next, these nuclei segmentation masks were used to estimate whole cell segmentation boundaries using the ‘skimage.morphology.binary_dilation’ function in the Python scikit-image package^105^. This function dilates nuclear segmentation boundaries by stochastically flipping pixels into the mask boundary with a probability equal to the fraction of positive neighboring pixels for 9 cycles. We then computed mean expression for each antibody across pixels within each whole cell segmentation boundary, which we define as the signal intensity 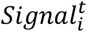 for cell *i and* target *t*.

##### Cell-level quality control

Since there is technical variation in CODEX staining and imaging quality, we applied multiple quality control filters to eliminate cells with atypical quality characteristics. First, we defined for cell *i* the signal sum Σ_*i*_, mean μ_*i*_, standard deviation σ_*i*_, and coefficient of variation CoV_*i*_ across the set of targets *T*, composed of DAPI + all antibodies in **Supplementary Table 3**.

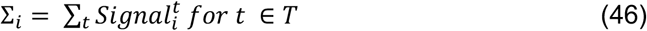

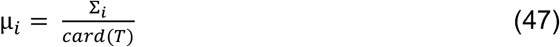

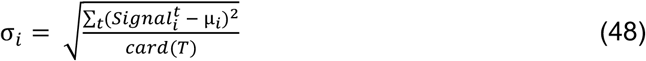

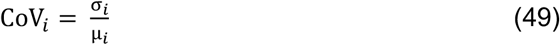

We then filtered cells for analysis by removing outliers for Σ_*i*_, CoV_*i*_, and 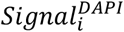 using manual thresholding of outlier thresholds and representative visual inspection to confirm low quality cells.

##### Annotation of cell types

Unsupervised clustering was performed by computing nearest neighbor distances (n_neighbors = 30) and Leiden clustering (resolution = 1) using the Python scanpy package^106,107^. Clustering was done simultaneously using all NSCLC mIF datasets. Clusters were annotated by a board-certified pathologist based on mean marker expression and manual inspection of raw mIF data. Following this, lymphocytes, macrophages, and indeterminate/mixed populations were subjected to sub-clustering and fine-level annotation. Finally, the initial annotations and fine annotations were merged to create a unified annotation of cell types across NSCLC mIF datasets (**Supplementary Table 4**).

##### Annotation of neighborhood types

Unsupervised identification of neighborhood clusters was performed by 1) defining local regions consisting of the 25 nearest cells in physical space and 2) clustering these regions into classes based on the frequencies of cell annotations they contain. This was done by defining the number of classes to be either 10 (‘neighborhoods’) or 30 (‘sub-neighborhoods’). Finally, classes were annotated by a board-certified pathologist based on the mean frequencies of their member cell annotations and by manual inspection of raw mIF data (**Supplementary Table 4**).

#### Markov Chain Monte Carlo (MCMC) simulation

##### CODEX mIF intensity normalization

Prior to using CODEX mIF intensities as an input to MCMC, we normalized signal intensities for (1) variation in local background and (2) variation in signal distribution between samples.

To correct for variation in local background, we divided each sample into 100 equally sized bins and used multi-Gaussian modeling for each target *t* ∈ *T* to identify the upper limits of that marker’s local null distribution. Let *i* and *j* represent the bin numbers in the *x* and *y* directions respectively. Then we denote *cells*_*i,j*_ as the set of cells in a given sample bin *(i,j)* and 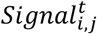 as the set of signal intensities for marker *t* for *cells*_*i,j*_. We used the ‘mclust’ R package to fit 2 Gaussians to 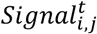 for all values of *i, j*, and *t*. Then we defined the upper bound 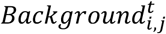 of the null distribution as the 95% percentile of that distribution for a given bin and marker. Finally, we subtracted 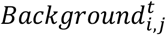 from 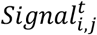 as a correction for local background signal variation.

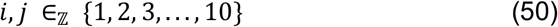

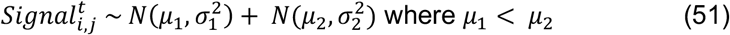

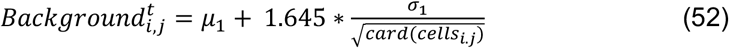

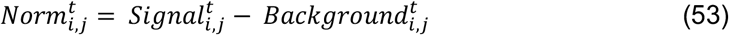

Cell-level normalized marker intensity data is contained within **Supplementary Table 4**.

##### MCMC data inputs

As depicted in **Fig. 6A**, there were four representations of the mIF data that were used as inputs to the MCMC simulation: 1) mean marker intensity, 2) cell type abundance, 3) neighborhood abundance, and marker SLAB. For each representation, mIF data coordinates were aligned to ST-seq spot coordinates (see ‘*Alignment of ST-seq and mIF data’*), and only the intensity data and annotations corresponding to cells located within ST-seq spot boundaries were considered. We found that amongst the six datasets with aligned ST-seq and mIF data, the mean number of mIF cell segmentations per ST-seq by dataset ranged from 15.2 cells/spot to 23.3 cells/spot (**Fig. S16**).

For mean marker intensity, we computed the mean intensity for each marker across all cells within a dataset. For cell type and neighborhood abundance, we computed the frequency of each cell type or neighborhood type annotation across all cells within a dataset. For marker SLAB, we first computed the sum of marker intensities for all cells in a spot. This resulted in a matrix of summed marker intensities where each spot in a dataset is arranged in rows and each mIF marker is arranged in columns. This matrix and the ST-seq spot coordinate matrix were used as inputs for TumorSPACE to compute SGs, differential marker abundance across SGs, and marker-wide SLAB scores as described in ‘*TumorSPACE: models and associated analysis*’. With each representation, the final matrix used as input to MCMC had dimensions of 6 samples x *f* features, where *f* is the number of features for that data representation.

##### MCMC algorithm

To perform empirical feature selection so that tumor spatial similarity defined by genome-wide SLAB could be compared to the four mIF data representations as described in **Fig. 6**, we applied a Markov Chain Monte Carlo (MCMC) approach. This was useful given the following properties of this task. First, this is a cardinality-contained optimization problem and therefore is NP-hard due to the discrete nature of feature selection. This requires solving the problem ‘in-shell’ for each cardinality ranging from one to the total number of features corresponding to a particular mIF representation. Furthermore, due to the lack of an analytical model that relates genome-wide SLAB to the mIF feature space, we set our objective (ℒ) as the correlations between SLAB properties (*D*^*SLAB*^) and equivalent properties produced by our feature set (*D*^*mIF*^). This correlation objective function is non-convex, meaning the search space possibly contains multiple local optima, thus rendering standard gradient-based optimization algorithms infeasible.

To address this, we created a Markov Chain Monte Carlo (MCMC) simulation framework as follows. For any cardinality ‘*k*’, the procedure begins by selecting a random subset of *k* features (*S*) referred to as ‘the seed’ out all possible *k*-membered subsets (*S*_*k*_) of total mIF-feature space ***F***_*mIF*_. From there, the MCMC algorithm enters a ‘high-entropy burn-in’ phase of broad exploratory sampling for 10,000 steps during which it probabilistically accepts steps in order to allow the system to escape local optima and survey a wide range of feature selection possibilities with a bias towards ones that increase the objective. Following this period, the algorithm transitions to a more focused ‘low-entropy convergence’ phase of 15,000 steps by starting with the optimized configuration from the burn-in phase and finetuning the feature selection by only accepting small changes that increase the objective.

Let *S*_*k*_ be the set of all k-membered subsets of mIF Features (*F*_*mIF*_), ℒ denote the correlation objective to be maximized, *S*^*^ denote the element of *S*_*k*_ that maximizes this objective, and *D*^*mIF*^(S) be the SLAB distances produced by a an element *S* of *S*_*k*_.

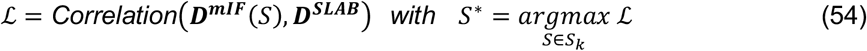

### Experimental method details

#### 10X Visium CytAssist spatial transcriptomics (ST-seq)

Tissue quality was determined by isolation of RNA from FFPE using the Qiagen RNeasy FFPE kit. Samples were then analyzed for tissue extraction quality using the Agilent 2100 bio-analyzer and Agilent RNA-6000 pico kit. For each sample, a DV200 score – the fraction of RNA fragments > 200 nucleotides in length – was calculated. Tissue quality for all samples was tested on unstained sections adjacent to the section used for ST-seq.

DLBCL samples were previously H&E stained. Imaging and coverslip removal were completed as described by 10X Protocol CG000518-Rev A and decrosslinking was performed according to 10X Protocol CG000520-Rev A^108,109^. NSCLC samples underwent deparaffinization, H&E staining, imaging, and decrosslinking according to CG000520-Rev B^110^. Sample imaging for all samples was performed using the Akoya Biosciences Vectra Polaris at 20X magnification.

We next performed the following steps as per either 10X Protocol CG000495-Rev A for the DLBCL samples or 10X Protocol CG000495-Rev E for the NSCLC samples^111,112^. First, samples underwent probe hybridization with Visium Human Transcriptome Probe Set v2.0, followed by probe ligation, and associated washes (**Supplementary Table 12**). Two native tissue slides and one Visium CytAssist 11 x 11 mm slide were then placed within the Visium CytAssist to enable RNA digestion, tissue removal, and transfer of ligated products onto the two fiducial frames of the Visium Slide. Next, we performed probe extension and elution off the Visium Slide, followed by pre-amplification and SPRIselect cleanup. For SPRIselect cleanup, DLBCL samples placed in only the ‘High’ position of the 10X magnetic separator, while NSCLC samples were placed in both ‘High’ and ‘Low’ positions according to CG000495-Rev E. To identify the optimal number of cycles for library amplification, we performed qPCR using Applied Biosciences QuantStudio 6 Pro as per CG000495-Rev E (**Supplementary Table 8**). For this step, we included 0.5 μl of carboxy-X-rhodamine (ROX) with the DLBCL samples and not with the NSCLC samples. Sample Index PCR was run using the sample-specific optimal number of cycles, followed by: cleanup, Agilent TapeStation QC, sequencing, and demultiplexing using Bcl2fastq. Sample sequencing was performed on a NovaSeq 6000 for DLBCL samples and a NovaSeqX for NSCLC samples. Sample-specific parameters and QC are listed in **Supplementary Table 8**. For DLBCL experiments, we used the Applied Biosystems Veriti 96 well thermocycler, while for NSCLC samples we used the Eppendorf Mastercycler X50a and X50I.

#### CODEX multiplexed immunofluorescence

##### Slide preparation

NSCLC samples were obtained as unstained slides mounted with 5 μm thickness formaldehyde-fixed, paraffin-embedded (FFPE) sections from the same patient biopsies as described in *‘Patient samples’*. Coverslips were coated with 0.1% poly-l-lysine solution prior to mounting tissue sections to enhance adherence. The prepared coverslips were washed and stored according to guidelines from the CODEX user manual^113^.

##### Antibody preparation

Custom conjugated antibodies were conjugated using the CODEX conjugation kit as per the CODEX user manual (**Supplementary Table 3**). Briefly, the antibody is (1) partially reduced to expose thiol ends of the antibody heavy chains, (2) conjugated with a CODEX barcode, (3) purified, and (4) added to Antibody Storage Solution for long-term stabilization. Subsequently, antibody conjugation is verified using sodium dodecyl sulfate-polyacrylamide gel electrophoresis and with QC staining.

##### Staining and data acquisition

Sample slides are stained following protocols in the CODEX User Manual. Briefly, samples are pretreated by heating at 60°C overnight, followed by deparaffinization, rehydration using ethanol washes, and antigen retrieval via immersion in Tris-EDTA pH 9.0 for 20 minutes. Samples are then blocked in staining buffer and incubated with the antibody cocktail for 3 hours at room temperature. After incubation, samples are washed and fixed following the CODEX User Manual. Data acquisition was performed using the PhenoCycler-Fusion 2.0 with a 20X objective, resulting in a resolution of 0.5 μm/pixel.

#### Patient tumor PD-L1 IHC

FFPE biopsy samples were probed for PD-L1 expression using a qualitative immunohistochemical assay with the Dako 22C3 antibody (Pharm Dx kit). PD-L1 expression was classified using the Tumor Proportion Score (TPS), which represents the percentage of viable tumor cells that show partial or complete membrane staining. Normal background histiocytes served as internal controls to ensure quality of the PD-L1 staining. Quantification was performed by a board-certified pathologist as part of routine clinical care.

#### Patient somatic mutation testing

The molecular profiles of the tumor biopsies were analyzed using Oncoplus or Oncoscreen, two Next Generation Sequencing (NGS) assays^114^. A description of patient mutation status can be found in **Supplementary Table 13**. Since the list of targeted genomic regions varied by the year in which testing was performed, a list of Oncoplus/Oncoscreen versions used for each patient as well as a list of the targeted genomic regions for each version can be found in **Supplementary Tables 13 and 14**, respectively.

For the Oncoplus analysis, DNA was isolated from the samples using the QIAamp DNA Blood Mini Kit (Qiagen), fragmented, and prepared into a sequencing library with patient-specific indexes (HTP Library Preparation Kit, Kapa Biosystems). Targeted genomic regions were enriched using a panel of biotinylated oligonucleotides (SeqCap EZ, Roche Nimblegen) supplemented with additional oligonucleotides (xGen Lockdown Probes, IDT). The enriched libraries were then sequenced on an Illumina HiSeq 2500 system, and the data was analyzed via bioinformatics pipelines against the hg19 (GRCh37) human genome reference sequence.

For Oncoscreen, DNA was isolated from formalin-fixed paraffin-embedded (FFPE) tumor tissue using the QIAamp DNA FFPE Tissue Kit (Qiagen). DNA was quantified using the Qubit fluorometric assay (Thermo Fisher Scientific) and a quantitative PCR assay (hgDNA Quantitation and QC kit, KAPA Biosystems). Targeted genomic regions were amplified using multiplex PCR (Thermo Fisher Scientific); PCR products were used to prepare NGS libraries with patient-specific adapter index sequences (HTP Library Preparation Kit, KAPA Biosystems). The enriched libraries were then sequenced on an Illumina MiSeq system, and the data was analyzed via bioinformatics pipelines against the hg19 (GRCh37) human genome reference sequence.

#### Patient tumor volume measurements

For measurement of tumor volume changes over time, computed tomography (CT) imaging reports were obtained for patients in the NSCLC as permitted by the IRBs referenced in ‘Patient samples’. For patients with measurable disease at the time of treatment start (denoted as month zero), the largest lesion was identified and labeled the ‘index lesion’. Changes in index lesions were collected when described in serial reports by a board-certified radiologist as part of routine clinical care.

### Subject details

#### Patient samples

Non-small cell lung cancer (NSCLC) patients were treated with immune checkpoint blockade therapy +/- chemotherapy at the University of Chicago Medical Center (Chicago, IL). All patients provided written informed consent for the collection and study of pre-treatment diagnostic tumor biopsy samples and for clinical outcomes including treatment regimen, treatment-related toxicities, and disease outcomes, as approved by the University of Chicago Institutional Review Board (IRB 9571 and IRB 24-0063). For the ST-seq analysis, 16 tumor samples were collected prior to therapy initiation, each from a separate patient. Inclusion criteria for these patients included (1) NSCLC stage IV patients either at initial presentation or as progression from previously treated early-stage disease, (2) biopsy of either the primary tumor or a metastatic tumor performed and stored within 6 months prior to treatment in the metastatic setting, (3) subsequent first line treatment with anti-PD1/anti-PD-L1 immune checkpoint blockade (ICB) with or without platinum-based chemotherapy. Exclusion criteria included (1) no prior therapy in the metastatic setting and (2) less than 2 doses of ICB therapy administered. We selected the first 16 patients that met these criteria and that had an available FFPE tumor biopsy block. From the archival block, a fresh 5 μm section was cut and placed on a standard slide for use in ST-seq protocols (see ‘*10X Visium spatial transcriptomics (ST-seq)*’). Progression was defined as time from the first dose of ICB until either radiographic or symptom-based evidence of disease progression. ICB regimen, ICB treatment duration, reason for ICB discontinuation, time to progression following ICB start, and time to death following ICB start are listed for all patients (**Supplementary Table 13**).

Diffuse large B-cell lymphoma (DLBCL) patients were treated at the University of Chicago Medical Center (Chicago, IL). All patients provided written informed consent for the collection and study of pre-treatment diagnostic tumor biopsy samples and for clinical outcomes including treatment regimen, treatment-related toxicities, and disease outcomes, as approved by the University of Chicago Institutional Review Board (IRB 13-1297). Each biopsy was reviewed by 2 hematopathologists for diagnostic confirmation. Biopsy slides were previously cut from FFPE sections and H&E stained for prior studies^115^.

## Supplementary Tables

Supplementary Table 1. Pan-tumor database properties

Supplementary Table 2. Pathologist annotation of pan-tumor H&E image data

Supplementary Table 3. CODEX Antibody Targets

Supplementary Table 4. CODEX normalized marker intensities and pathologist annotations

Supplementary Table 5. Differential spatial lability genes amongst pan-tumor SLAB classes

Supplementary Table 6. Differential spatial lability pathways amongst pan-tumor SLAB classes

Supplementary Table 7. Bulk gene sets for prediction of NSCLC ICB response

Supplementary Table 8. ST-seq QC Metrics

Supplementary Table 9. Schema for pathologist annotation of H&E image data.

Supplementary Table 10. SpaCET input cancer types

Supplementary Table 11. Reactome pathway list

Supplementary Table 12. Visium Human Transcriptome Probe Set v2.0 - Probe Set Reference CSV file

Supplementary Table 13. NSCLC Clinical Metadata (oncoplus version)

Supplementary Table 14. Oncoplus/Oncoscreen Gene Panels

## Acknowledgements

We thank D. Pincus, M. Mani, M. Lingen, D. Zemmour, I. Moskowitz, and R. Ranganathan for helpful discussions. We thank the genomics core and human immunologic monitoring core facilities at the University of Chicago for their aid in sequencing and imaging our ST-seq samples. This work was supported by the Duchossois Family Institute, the Department of Pathology, and the Center for the Physics of Evolving Systems at the University of Chicago. Funding for V.B. was provided by a T32 NIH training grant within the Department of Medicine, Section of Hematology/Oncology at the University of Chicago.

## Author contributions

A.G. is a board-certified pathologist who performed all pathologist annotations and performed data analysis. H.G. performed all ST-seq data collection including preparing samples and running the 10X Visium platform. U.P. provided critical conceptual guidance in writing the manuscript and conducted portions of analysis related to Figs. 3 and 5. A.D.L., A.E., A.P., C.M.B. aided in data collection of the NSCLC samples. A.D.L. aided in the writing of the Methods section. B.P. provided critical conceptual input and technical support in performing Monte Carlo simulations. B.A.D. provided critical conceptual guidance as well as technical support in writing of the code used for TumorSPACE. A.H. is a board-certified pathologist who supervised pathologist annotations and assisted with interpretation of annotations. C.M.B. provided critical feedback for our manuscript. J.K. provided diffuse large B cell lymphoma samples for ST-seq and conceptual guidance. M.C.G. leads the lung cancer biobank at the University of Chicago, established the IRB required for collecting ST-seq data on the NSCLC samples, and provided critical feedback for our manuscript. V.B. performed all analysis, wrote all code, and coordinated all data collection efforts. V.B. and A.S.R. conceived of the project and A.S.R. provided supervision for all aspects of analysis and data collection. V.B. and A.S.R. wrote the paper.

## Competing interests

A.S.R. is a founder of Sparsity, Inc. V.B. is a co-founder of Sparsity, Inc.

## Data and materials availability

All data relevant to our manuscript can be found within associated Supplementary Tables. All ST-seq data related to the cohort of DLBCL and NSCLC patients are available at https://www.ncbi.nlm.nih.gov/geo/query/acc.cgi?acc=GSE292299. All pathologist-annotated H&E images corresponding to our pan-tumor database are available at 10.5281/zenodo.16856762.

## Code availability

All TumorSPACE code, along with documentation and step-wise instructions, will be available for download via github repository upon publication of our manuscript at https://github.com/aramanlab/TumorSPACE.jl. All code for producing the figures in this paper from source code will be available for download via github repository upon publication of our manuscript at https://github.com/aramanlab/Behera_etal_2025.

For the stage of review, any raw data and code repositories can be accessed as follows:

ST-seq data for GSE2922299:

1. Go to https://www.ncbi.nlm.nih.gov/geo/query/acc.cgi?acc=GSE292299
2. Enter token svsluumozfkxlkb into the box

Pathologist-annotated H&E data: https://zenodo.org/records/16856762?token=eyJhbGciOiJIUzUxMiIsImlhdCI6MTc1NTI2OTU1OCwiZXhwIjoxNzY3MTM5MTk5fQ.eyJpZCI6IjU4ODBhNmFhLWNmZTMtNGI3Ny1hMTkwLTg3ZTU3NjJhMTM2YSIsImRhdGEiOnt9LCJyYW5kb20iOiI5NjYxMTNiODIzN2FkNGZkNDQ1MjU3YTljZWVlNDMxOSJ9.Njxe6tjN7MPsRO8JHuF2rotE_aCIC-YjssqxSiASAFFbPaNAJRhG21M96i_5ZLmEtRzYJ9K80WratYjvNU9Qyw

TumorSPACE repo:

1. Open a terminal
2. Navigate to the folder with the private key (‘reviewer_deploykey_TumorSPACE)
3. Enter the following to clone the repo: GIT_SSH_COMMAND=“ssh -i reviewer_deploykey_TumorSPACE -o StrictHostKeyChecking=no” git clone git@github.com:aramanlab/TumorSPACE.jl.git
  a. If this command results in a warning such as ‘Permissions 0644 for ‘reviewer_deploykey_TumorSPACE’ are too open’, this is because the key file permission have changed from 600 to 644 during download. Set them back to 600 by entering the following: chmod 600 reviewer_deploykey_TumorSPACE
4. Within the repo folder, open the Installation instructions in docs/build/installation/index.html. Follow these for package installation and to run the tutorial.
5. Alternatively, can unzip the TumorSPACE.jl.zip repo directly.
6. To view the documentation as a live website, do the following:
  a. Navigate to TumorSPACE.jl/docs/build/
  b. Enter the following in the command line: python3 -m http.server 8000
  c. Open a browser and go to: http://localhost:8000

Paper repo (for reproducing figures):

1. Open a terminal
2. Navigate to the folder with the private key (‘reviewer_deploykey_Behera_etal_2025)
3. Enter the following to clone the repo: GIT_SSH_COMMAND=“ssh -i reviewer_deploykey_Behera_etal_2025 -o StrictHostKeyChecking=no” git clone git@github.com:aramanlab/Behera_etal_2025.git
  a. If this command results in a warning such as ‘Permissions 0644 for ‘reviewer_deploykey_Behera_etal_2025’ are too open’, this is because the key file permission have changed from 600 to 644 during download. Set them back to 600 by entering the following: chmod 600 reviewer_deploykey_ Behera_etal_2025
4. To view the figure generation as a live website, do the following:
  a. Navigate to Behera_etal_2025/docs/
  b. Enter the following in the command line: python3 -m http.server 8001
  c. Open a browser and go to: http://localhost:8001

## Ethics Declaration

Patents (63/572,XXX) related to this research have been filed by the University of Chicago with V.B. and A.S.R. as inventors.

---

**Figure S1.**
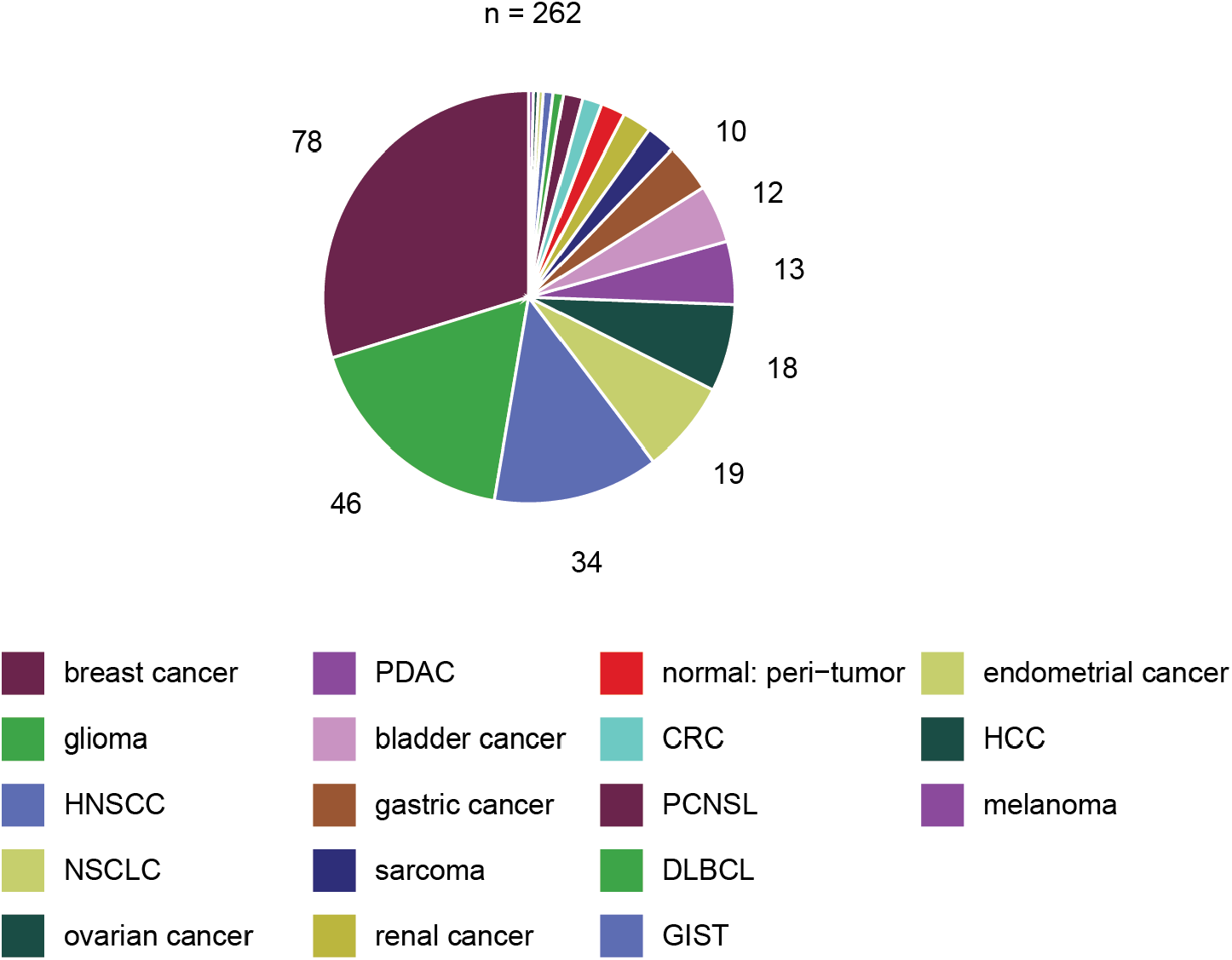
Description of our tumor database. Number of datasets for each tumor type (color key) is delineated in the pie graph.

**Figure S2.**
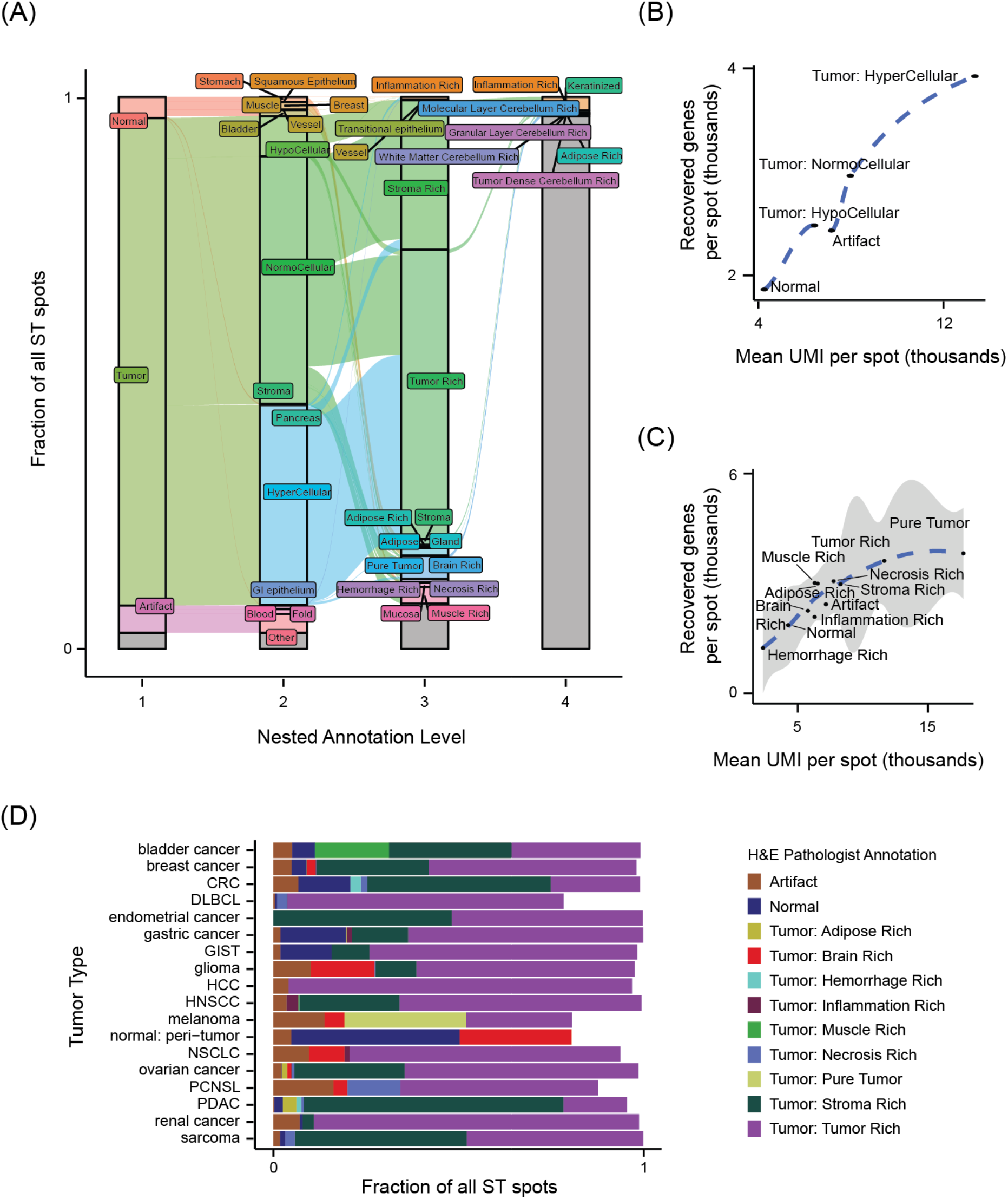
**(A)** Alluvial plot depicting pathologist annotation of ST spots for all pan-tumor datasets (y-axis) into a nested four-level hierarchy (x-axis). **(B,C)** Mean UMI (in thousands) per spot (x-axis) versus mean number of recovered genes (in thousands) per spot (y-axis) by tumor cellularity (B) or tumor region subtype (C). **(D)** Bar plot depicting the fraction of ST spots, stratified by tumor type (y-axis), classified into various pathologist annotations based on the paired H&E image (x-axis).

**Figure S3.**
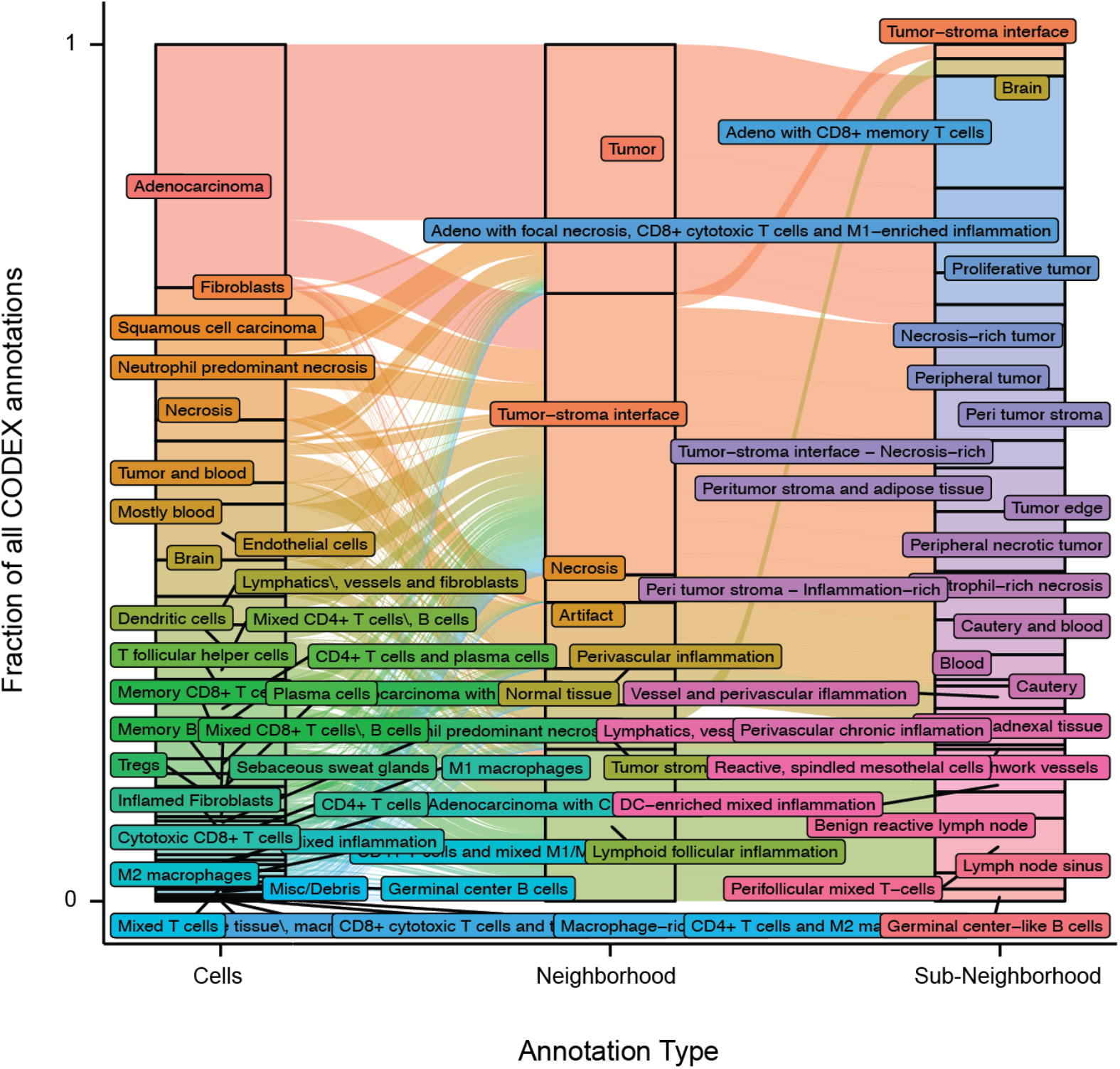
Alluvial plot depicting pathologist assignment of mIF cell segmentations into cell types (left), neighborhood types (middle), and sub-neighborhood types (right).

**Figure S4.**
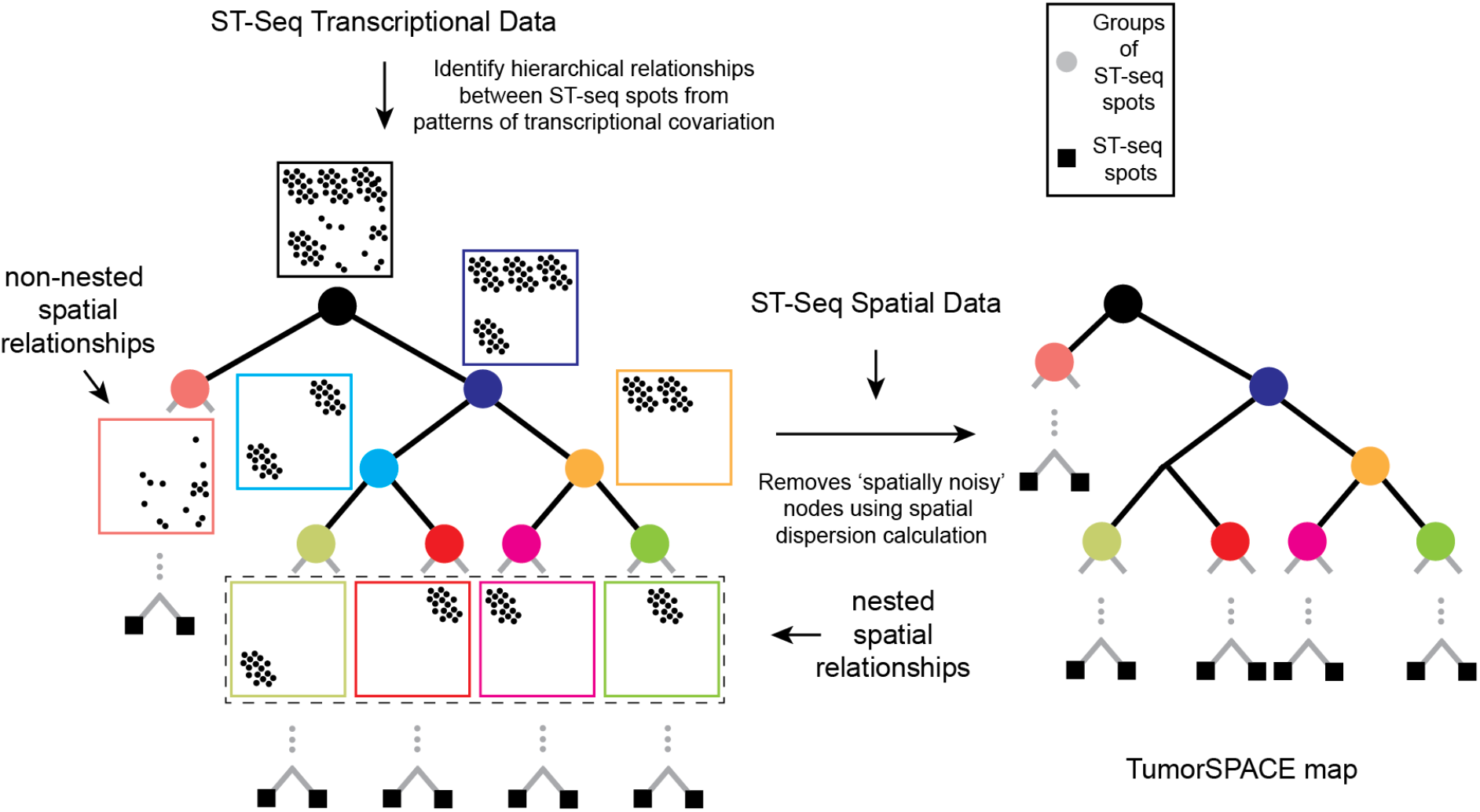
Workflow for generating a TumorSPACE map involves first identifying hierarchical relationships between ST-seq spots using transcriptional data alone (left) and then performing spatial de-noising by removing tree nodes with high spatial dispersion values (right) (Methods). These maps can capture both spatially nested and spatially non-nested spot relationships. Grey lines at the bottom of each branch point indicate that trees continue until terminating at the individual ST-seq spots (black squares).

**Figure S5.**
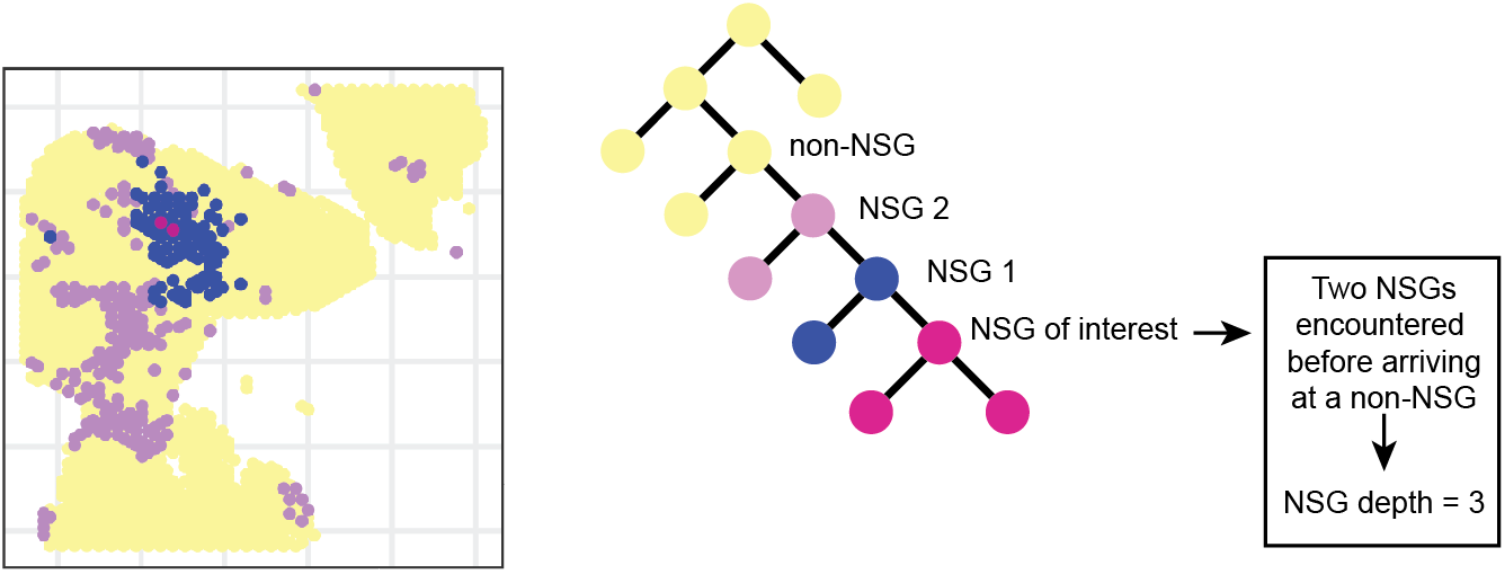
Description of how NSG depth is calculated for an example set of SGs.

**Figure S6.**
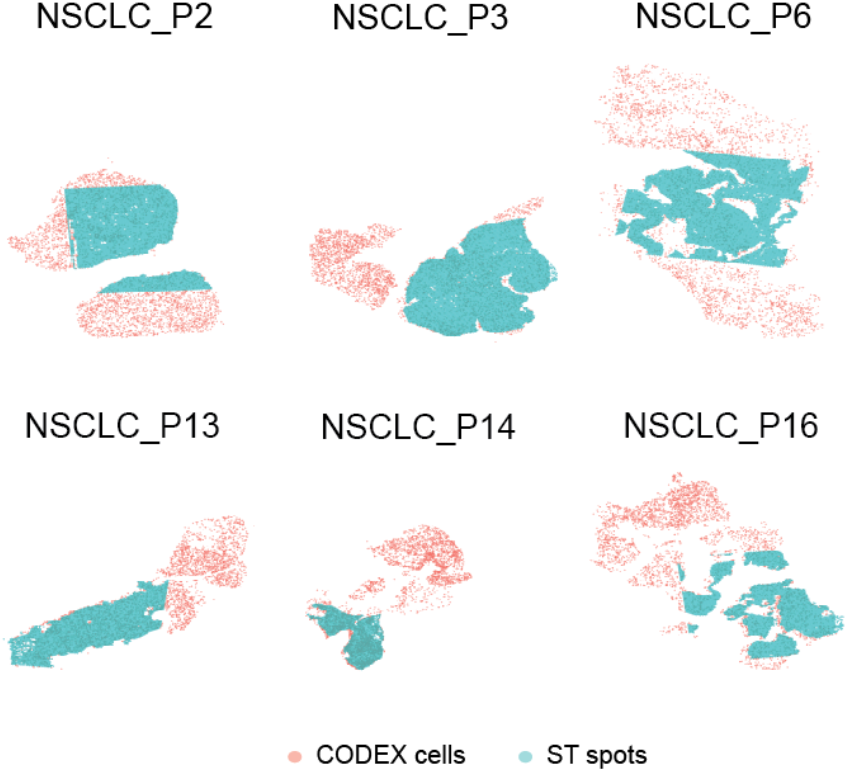
Alignment of ST spots (blue) to mIF cell segmentations (red) between serial tissue sections. Facets indicate distinct NSCLC patient biopsies.

**Figure S7.**
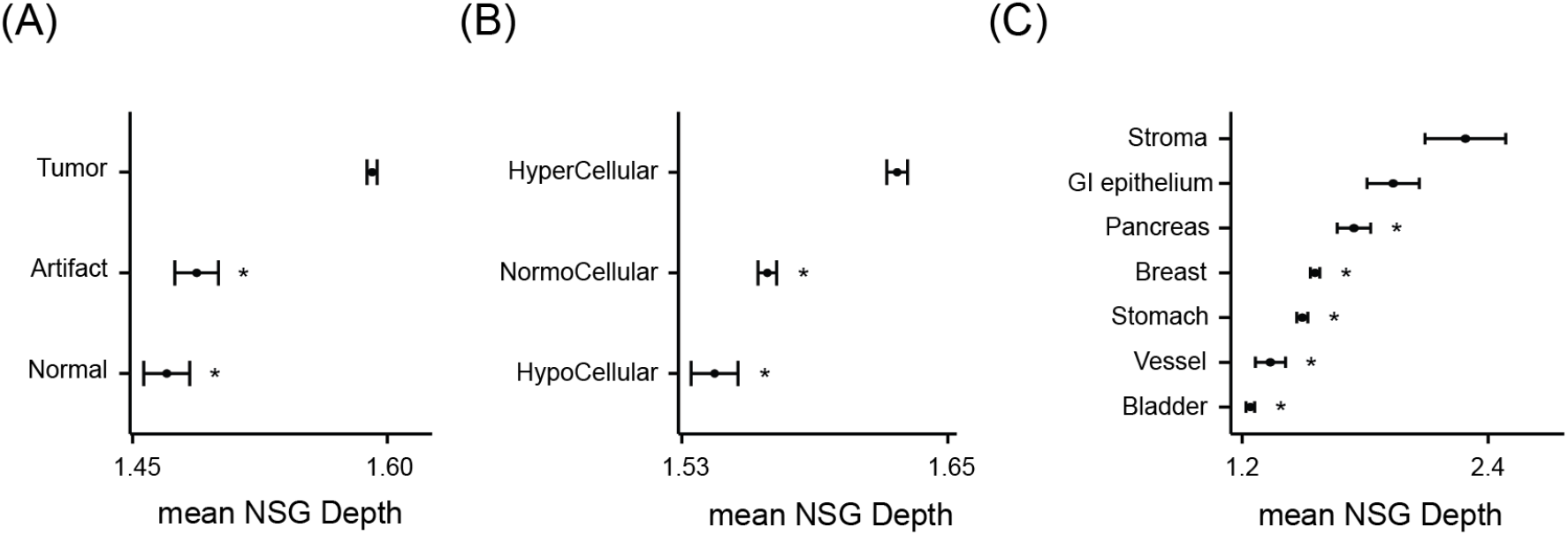
**(A - C)** Mean NSG depth of spots (x-axis) grouped by broad tissue type (A), tumor cellularity (B), or normal tissue sub-type (C), as annotated by a pathologist using H&E images. Non-NSGs are assigned a depth of 0. Error bars reflect the standard error of the mean. *Wilcoxon p-value < 0.05.

**Figure S8.**
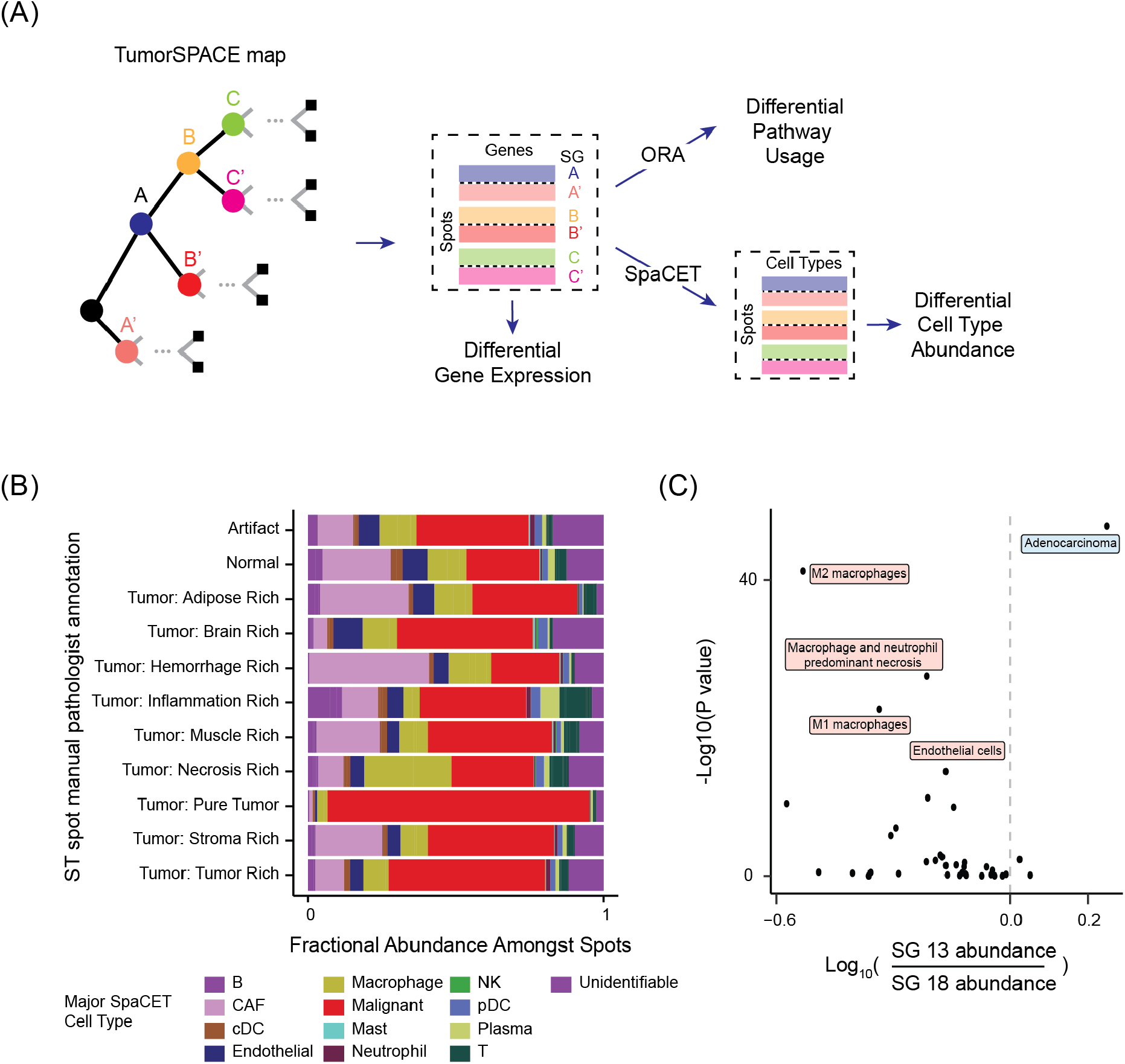
**(A)** Using TumorSPACE models to conduct differential analysis of gene expression, pathway usage with over-representation analysis (ORA), and cell type differences using SpaCET for spot deconvolution into cells. SGs are labeled as A, A’, B, B’, C, and C’. **(B)** Bar chart depicting the fraction of H&E-annotated ST spots (x-axis) deconvoluted into SpaCET major cell types (colors), stratified by pathologist annotation of tumor sub-type (y-axis). **(C)** Ratio in mean fraction (log-10 scale) of pathologist-annotated cell types from mIF data amongst cells aligned to SG13 vs SG18 (x-axis) vs log-transformed p-value of difference (y-axis). The most significant cell types enriched in either SG13 (blue) or SG18 (red) are labeled.

**Figure S9.**
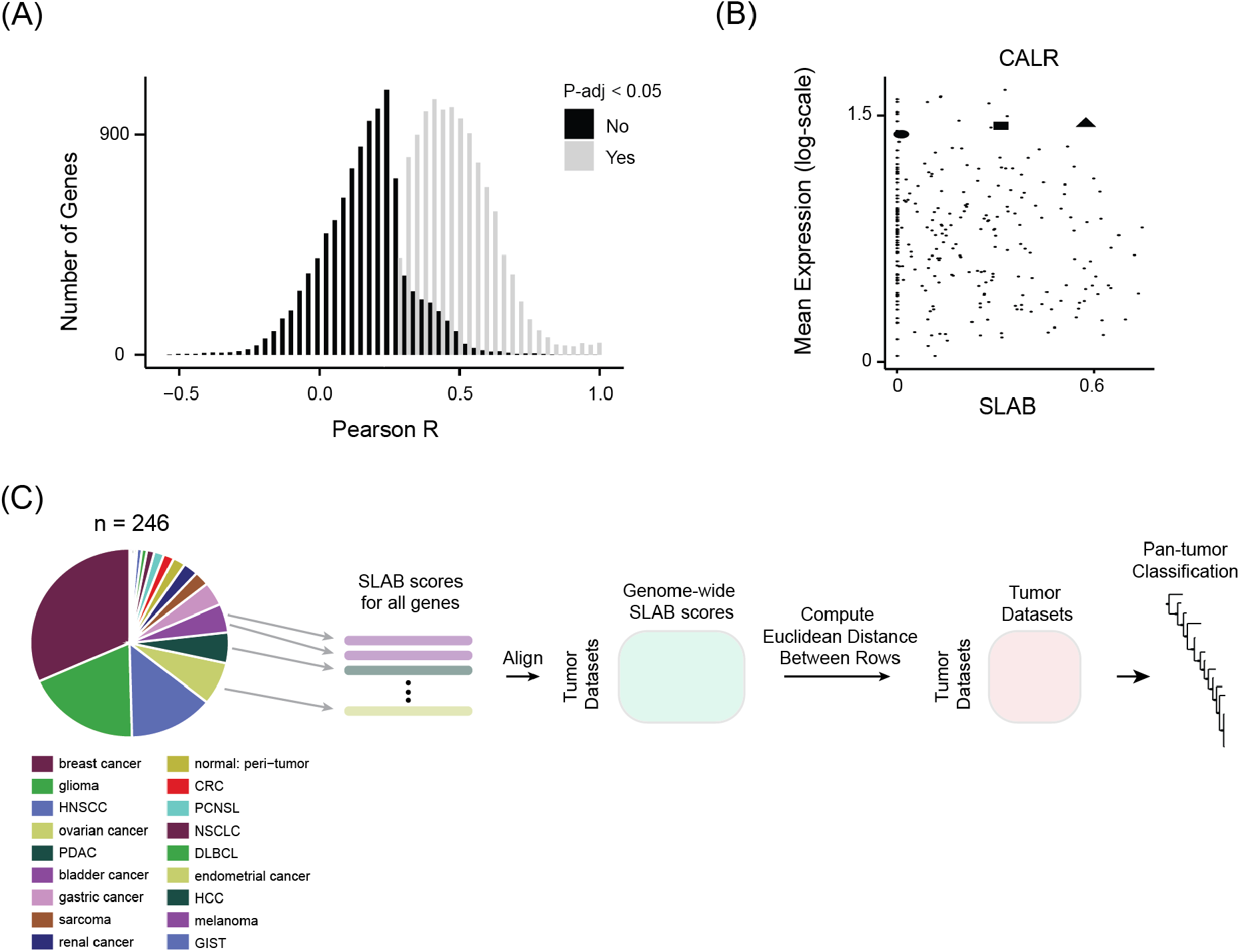
**(A)** Histogram of correlations (Pearson R) between bulk gene expression and SLAB scores across all tumors in our pan-tumor database for all genes. Genes are stratified by whether correlation was statistically significant (grey) or not (black) compared to an empirical null distribution. **(B)** Expression of CALR averaged across all spots in each tumor (y-axis) versus CALR SLAB score (x-axis). Each dot is a tumor in our database. Three tumors (black circle, square, and triangle) are highlighted that harbor the same mean CALR expression but varying SLAB scores. **(C)** Workflow for pan-tumor classification by SLAB scores. First, SLAB scores are computed for each gene for all tumors. This creates a genome-wide profile of SLAB scores for each ST-seq dataset. The datasets are aligned by their genome-wide SLAB profiles creating a matrix where rows are ST-seq datasets, columns are genes, and each entry is the SLAB score for a gene in an ST-seq dataset. Euclidean distance based on genome-wide SLAB scores is computed for all pairs of ST-seq datasets. Hierarchical clustering of pairwise SLAB-based distance results in a pan-tumor classification where tumors that are close together share a similar genome-wide SLAB profile.

**Figure S10.**
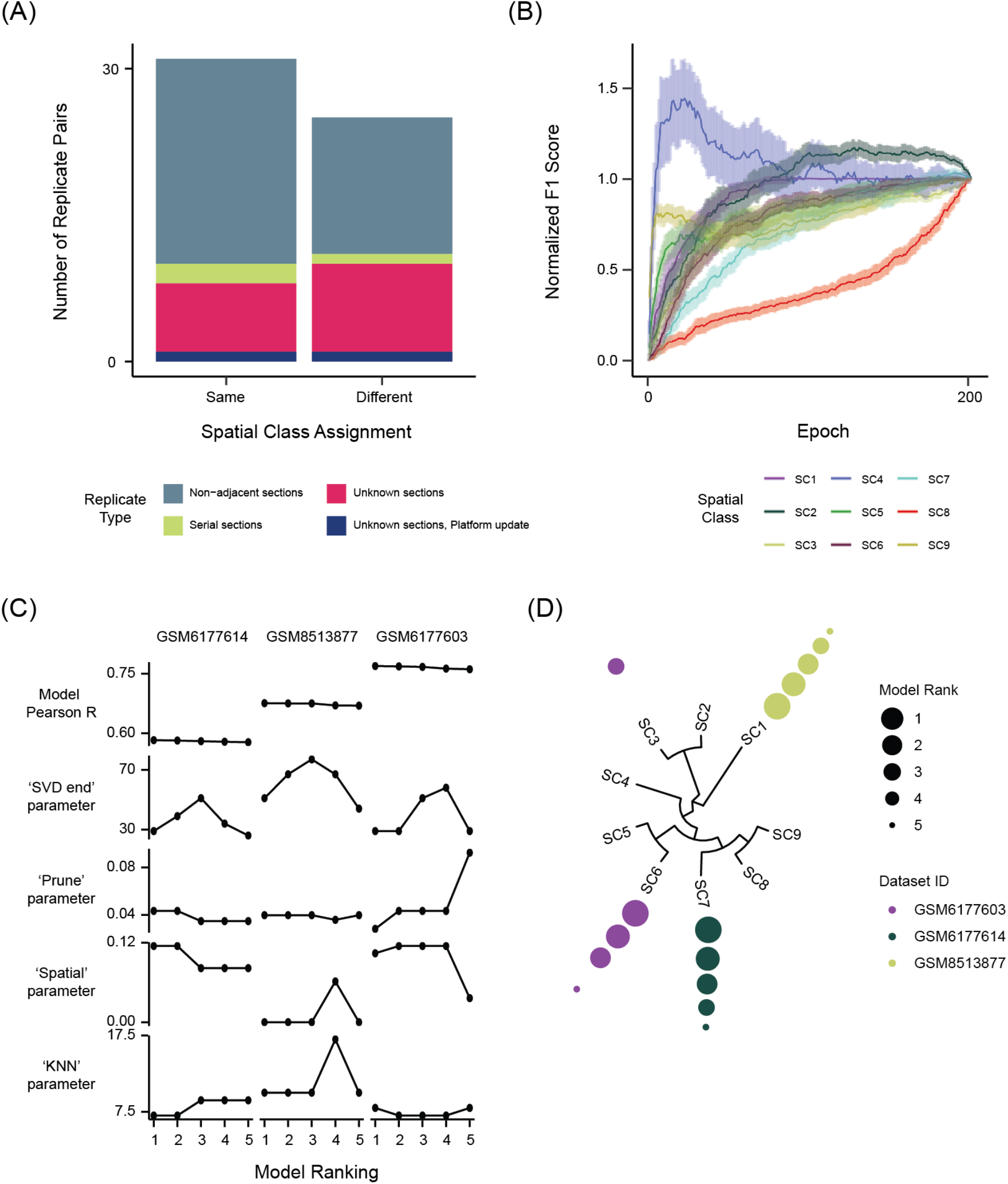
**(A)** Comparison of Spatial Class assignment (x-axis) across replicate pairs (color key). **(B)** Simulated construction of the pan-tumor latent space using a randomly selected order in which datasets are incorporated. The incorporation of all tumor sections corresponding to a single unique tumor constitutes one epoch. At each epoch, all datasets are projected into the simulated latent space. Plot depicts mean F1 score (y-axis) across 100 simulations for each Spatial Class across simulation epochs (x-axis). **(C)** Performance (Model Pearson R) and hyperparameter settings (vertical facets) for the five top-ranking TumorSPACE models (x-axis) across three representative datasets (horizontal facets). **(D)** Projection of top-ranking models (points, where size indicates model rank) into the pan-tumor classification for all three datasets (colorkey).

**Figure S11.**
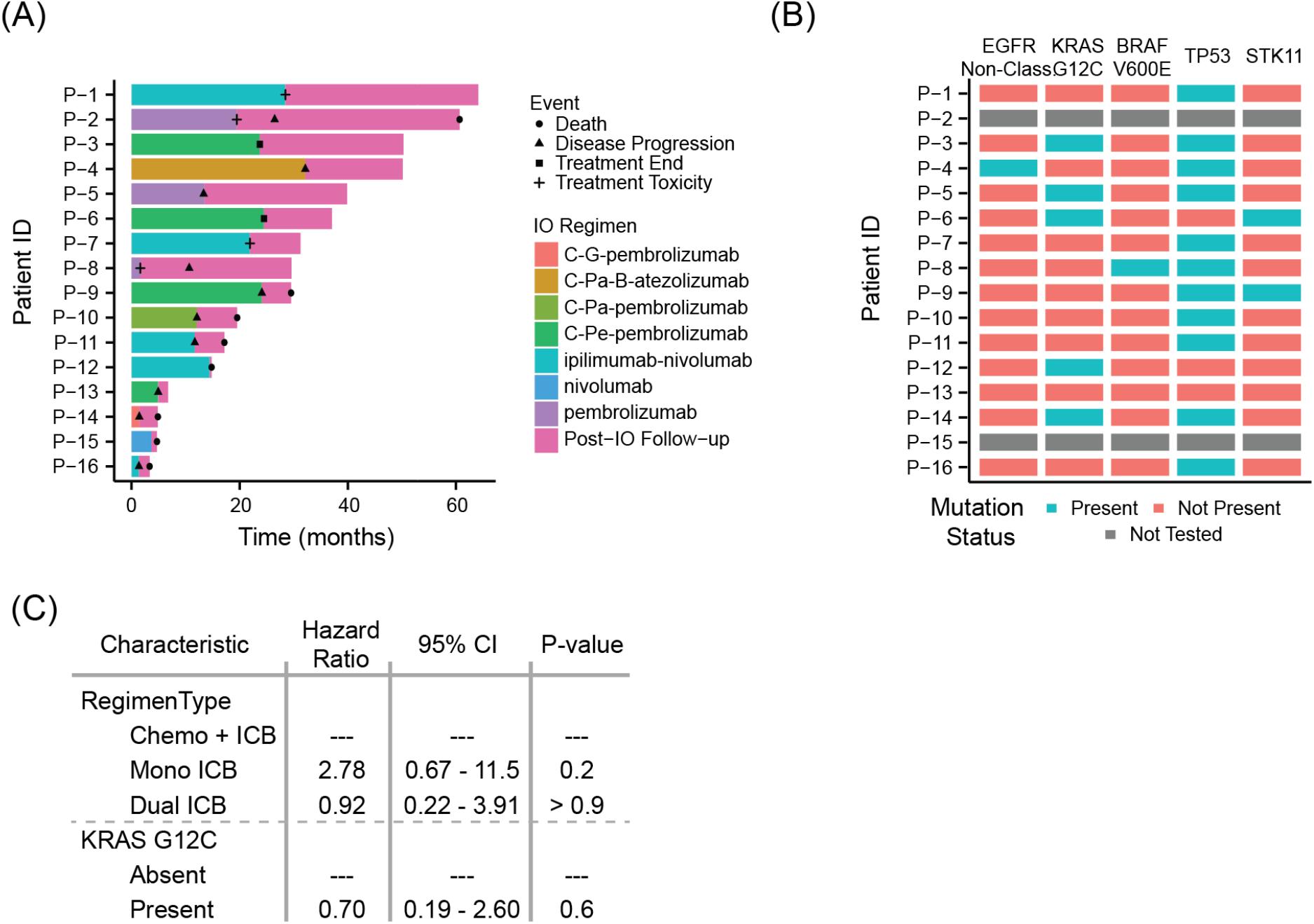
**(A)** Swimmer plot illustrating patient treatment courses starting when patients began frontline immunotherapy treatment in the metastatic NSCLC setting. Colors indicate immunotherapy (IO)/chemo-IO regimen, shapes indicate significant events. Chemotherapies are abbreviated as follows: C = carboplatin, G = gemcitabine, Pa = paclitaxel, B = bevacizumab, Pe = pemetrexed. IO therapies include anti-PD1 (pembrolizumab, nivolumab), anti-PD-L1 (atezolizumab), and anti-CTLA-4 (ipilimumab) therapies. **(B)** Mutation status for clinically relevant mutations amongst the 16-patient cohort at the time of pre-treatment diagnostic biopsy. **(C)** Univariate analysis between (i) ICB regimen type, and (ii) KRAS G12C status and progression-free survival (PFS).

**Figure S12.**
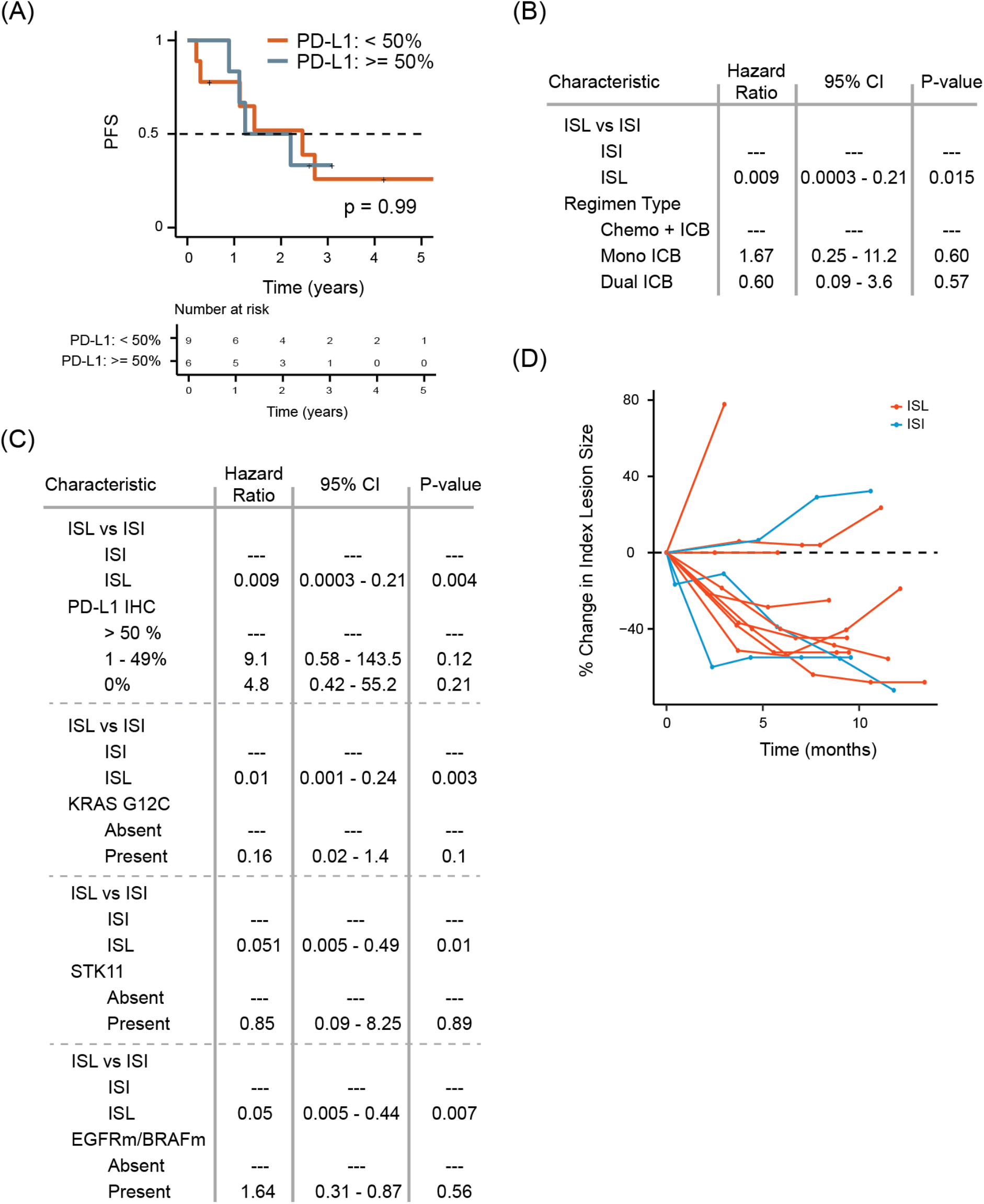
**(A)** Kaplan-Meier curve for PFS (y-axis) in years (x-axis) stratified by PD-L1 status using a binary threshold at 50%. **(B,C)** Multivariate PFS analysis of ISL/ISI with treatment regimen type (B) or somatic mutation status (C). **(D)** Spider plot depicting percent change in index tumor lesion volume using serial computed tomography (CT) scans (y-axis) following treatment start (x-axis). Each line describes a single patient classified as either ISL (blue) or ISI (red), and each point on a line indicates a CT scan measurement at that time.

**Figure S13.**
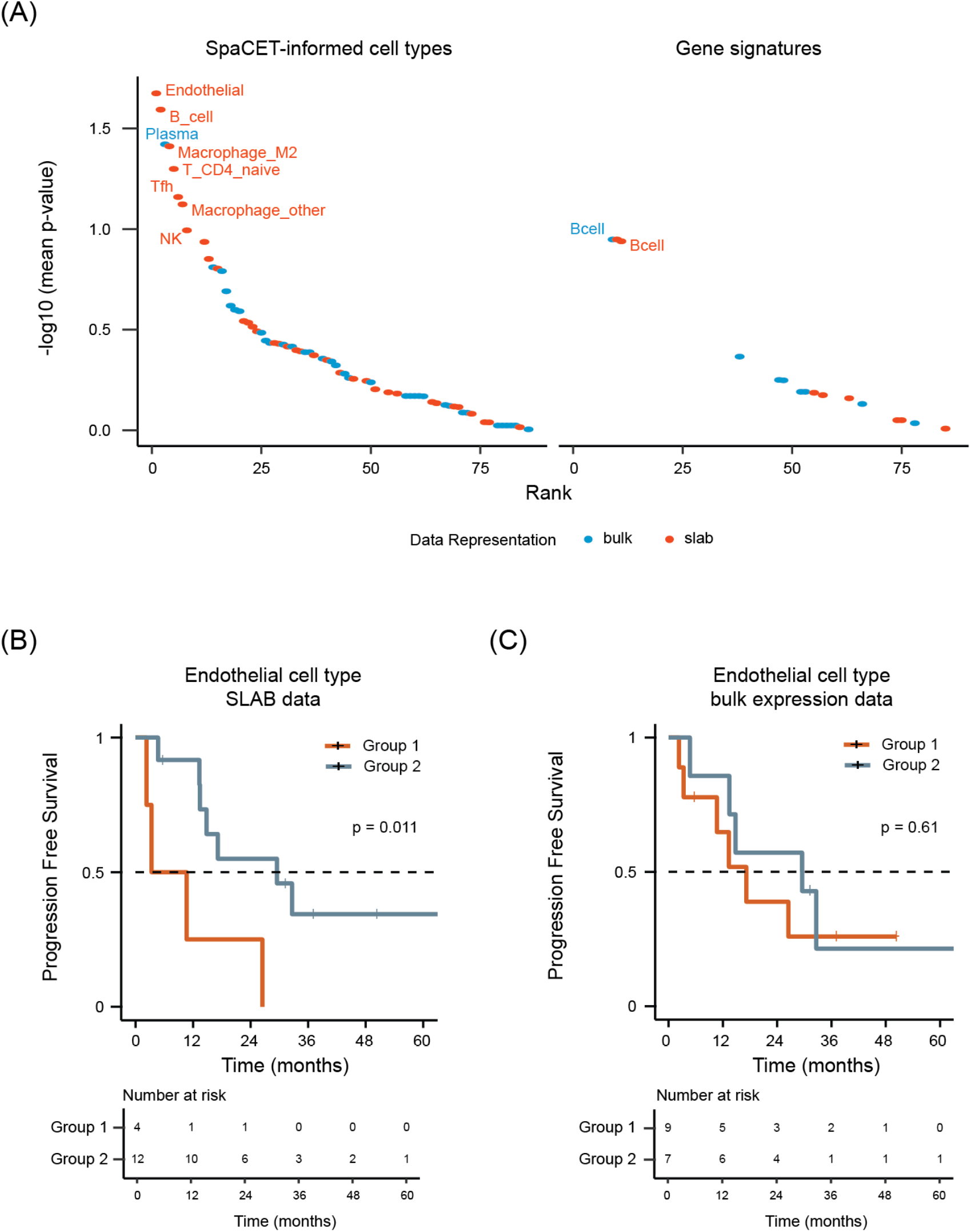
**(A)** Mean p-values (log10-transformed, y-axis) of log rank statistical tests for estimating PFS using SpaCET-inferred cell type abundance (left panel) or published bulk expression gene sets (right panel). **(B,C)** Kaplan-Meier curve for PFS (y-axis) when NSCLC patients are stratified by endothelial cell abundance using SLAB **(B)** or bulk abundance **(C)**.

**Figure S14.**
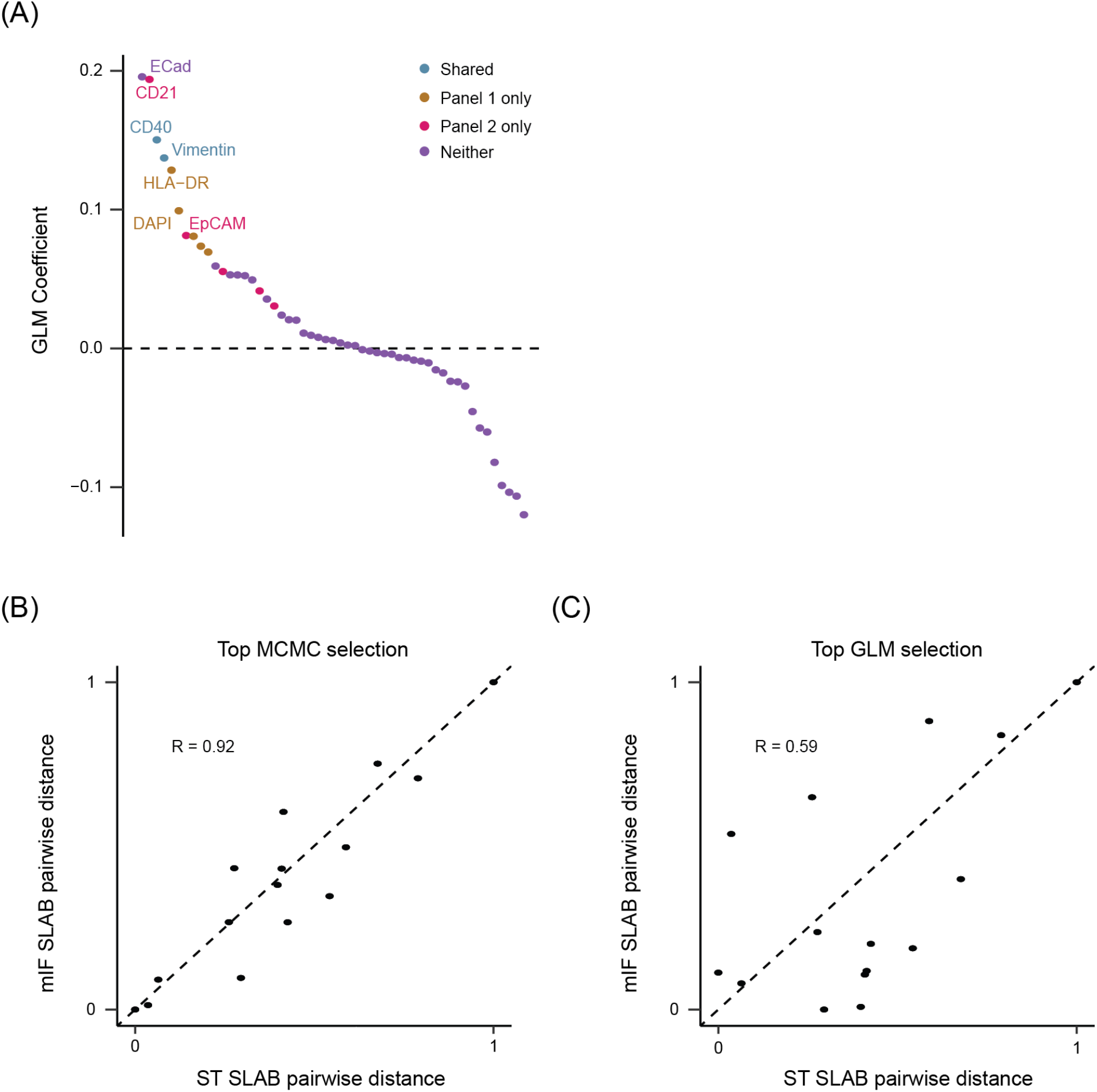
**(A)** Generalized linear model (GLM) regression coefficients (y-axis) of marker SLAB features (x-axis) for estimating sample similarities from genome-wide SLAB. MCMC simulation data from all panel sizes were used. Marker colors indicate membership in Panel 1, Panel 2, both (shared), or neither (see **Fig. 6B** for definition of panels with respect to constituent markers). **(B,C)** Normalized Euclidean distance between pairs of tumors based on mIF SLAB (y-axis) where markers were either a 7-member panel derived from MCMC **(B)** or GLM **(C)** frameworks versus normalized Euclidean distance between pairs of tumors based on genome-wide SLAB profiles (x-axis).

**Figure S15.**
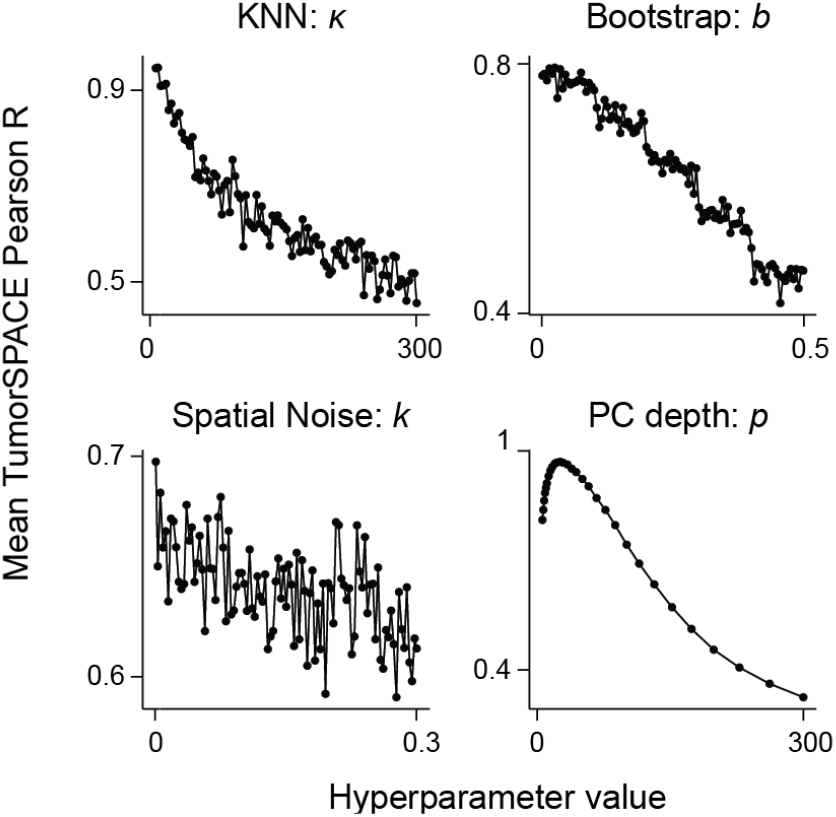
Mean TumorSPACE Pearson R across all pan-tumor models for particular hyperparameter values: KNN κ(top left), bootstrap confidence *b* (top right), spatial noise *k* (bottom left), and PC depth *p* (bottom right).

**Figure S16.**
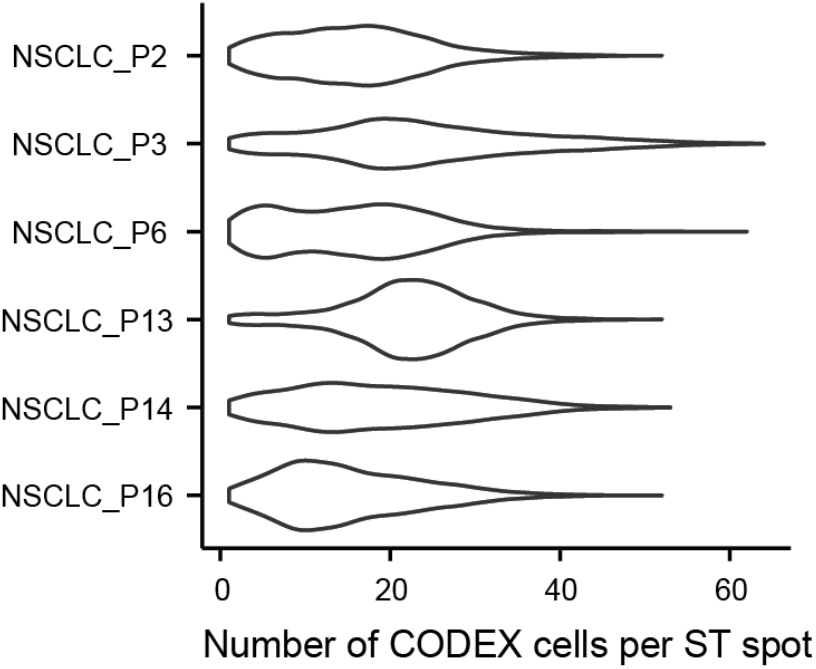
Violin plot depicting the number of mIF cell segmentations per ST spot (x-axis) for NSCLC samples (y-axis).

## Notes

https://github.com/aramanlab/TumorSPACE.jl

### Competing Interest Statement

The authors have declared no competing interest.

### Summary of Updates

This version has been updated to reflect revisions performed during the review process.

